# 3D SYNAPTIC ORGANIZATION OF THE RAT CA1 AND ALTERATIONS INDUCED BY COCAINE SELF-ADMINISTRATION

**DOI:** 10.1101/2020.04.23.050575

**Authors:** L. Blazquez-Llorca, M. Miguéns, M. Montero-Crespo, A. Selvas, J. Gonzalez-Soriano, E. Ambrosio, J. DeFelipe

## Abstract

The hippocampus plays a key role in contextual conditioning and has been proposed as an important component of the cocaine addiction brain circuit. To gain knowledge about cocaine-induced alterations in this circuit, we used Focused Ion Beam milling/Scanning Electron Microscopy (FIB/SEM) to reveal and quantify the 3D synaptic organization of the *stratum radiatum* of rat CA1, under normal circumstances and after cocaine-self administration (SA). Most synapses are asymmetric (excitatory), macular-shaped, and in contact with spine heads. After cocaine-SA, the size and complexity of both asymmetric and symmetric (inhibitory) synapses increased but no changes were observed in the synaptic density.

This work constitutes the first detailed report on the 3D synaptic organization in the *stratum radiatum* of the CA1 field of cocaine-SA rats. Our data contribute to the elucidation of the normal and altered synaptic organization of the hippocampus, which is crucial for better understanding the neurobiological mechanisms underlying cocaine addiction.

## INTRODUCTION

The mechanisms underlying the causes of drug addiction are not entirely known, but it is thought that the processes of learning and memory are essential and that the hippocampus plays a key role in contextual conditioning and cocaine addiction^1,2^. In fact, addiction has been considered as a type of non-adaptive learning^3^. In this regard, the hypothesis that drugs and natural enhancers act on similar brain systems^4,5^ is becoming more relevant. Numerous studies have found that a variety of drugs of abuse produce several morphological modifications of the pyramidal neurons, including the dendritic spines, which represent the main postsynaptic target of excitatory glutamatergic synapses of the cerebral cortex^6^. These changes are found in different brain regions including the frontal cortex, hippocampus and the nucleus accumbens (e.g.,^7,8,9,10,11,12,13,14^). Previous studies in our laboratory have shown that cocaine-induced changes in the hippocampus depend on the rat strain; specifically, an increase in the dendritic spine density and the proportion of larger spines after cocaine self-administration (SA) in Lewis rats^13,14^. Several global and local processes, involving dendritic spine and synaptic re-organization (structural plasticity) occurring in a defined temporal manner, have been proposed to provide a mechanism for long-term storage of memory traces upon learning that influences behavior^15,16^. For this reason, dendritic spine changes, among other structural changes, might explain functional alterations —such as facilitation of long-term potentiation (LTP) or impaired LTP depotentiation in the hippocampus— as well as behavioral alterations after cocaine-SA^17,18,19,20,21,22^. Therefore, changes in hippocampal dendritic spines seem to play a key role in cocaine addiction.

However, it has been well established via electron microscopy that dendritic shafts also form synapses and not all dendritic spines represent postsynaptic targets^23,24,25^. Thus, electron microscopy with serial section reconstruction is the gold standard method for studying the normal synaptic organization and its possible alterations. Here, we used Focused Ion Beam/Scanning Electron Microscopy (FIB/SEM) technology to perform a three-dimensional (3D) analysis of the synaptic organization in the neuropil in the *stratum radiatum* of the CA1 region of Lewis rats — an inbred rat strain that is prone to the reinforcing effects of drugs of abuse^26^. This hippocampal layer receives most afferents from the Schaffer collaterals of the CA3 pyramidal neurons — a synaptic connection that has a key role in the learning-induced synaptic potentiation of the hippocampus^27^. We first studied the normal (control) *stratum radiatum* and then compared the results with that obtained in the *stratum radiatum* of rats after intravenous cocaine-SA. This drug consumption model is considered the best for simulating the pattern of consumption in humans^28^. Specifically, we studied a wide variety of synaptic structural parameters including the synaptic density and spatial distribution; proportions of synapses; postsynaptic targets; as well as the shape and size of the synaptic junctions. This work constitutes the first detailed report of the 3D synaptic organization in the *stratum radiatum* of the hippocampal CA1 field of cocaine-SA rats. Our data contribute to the elucidation of the normal and altered synaptic organization of the rat hippocampus, which is crucial for better understanding the neurobiological mechanisms underlying cocaine addiction.

## RESULTS

We have analyzed the synaptic organization of the *stratum radiatum* of the CA1 hippocampal region of control Lewis rats (saline group) in serial sections and evaluated possible differences in this synaptic organization after cocaine-SA (**Figures 1, 2, 3 and Supplementary figure 1**).

**Figure 1.**
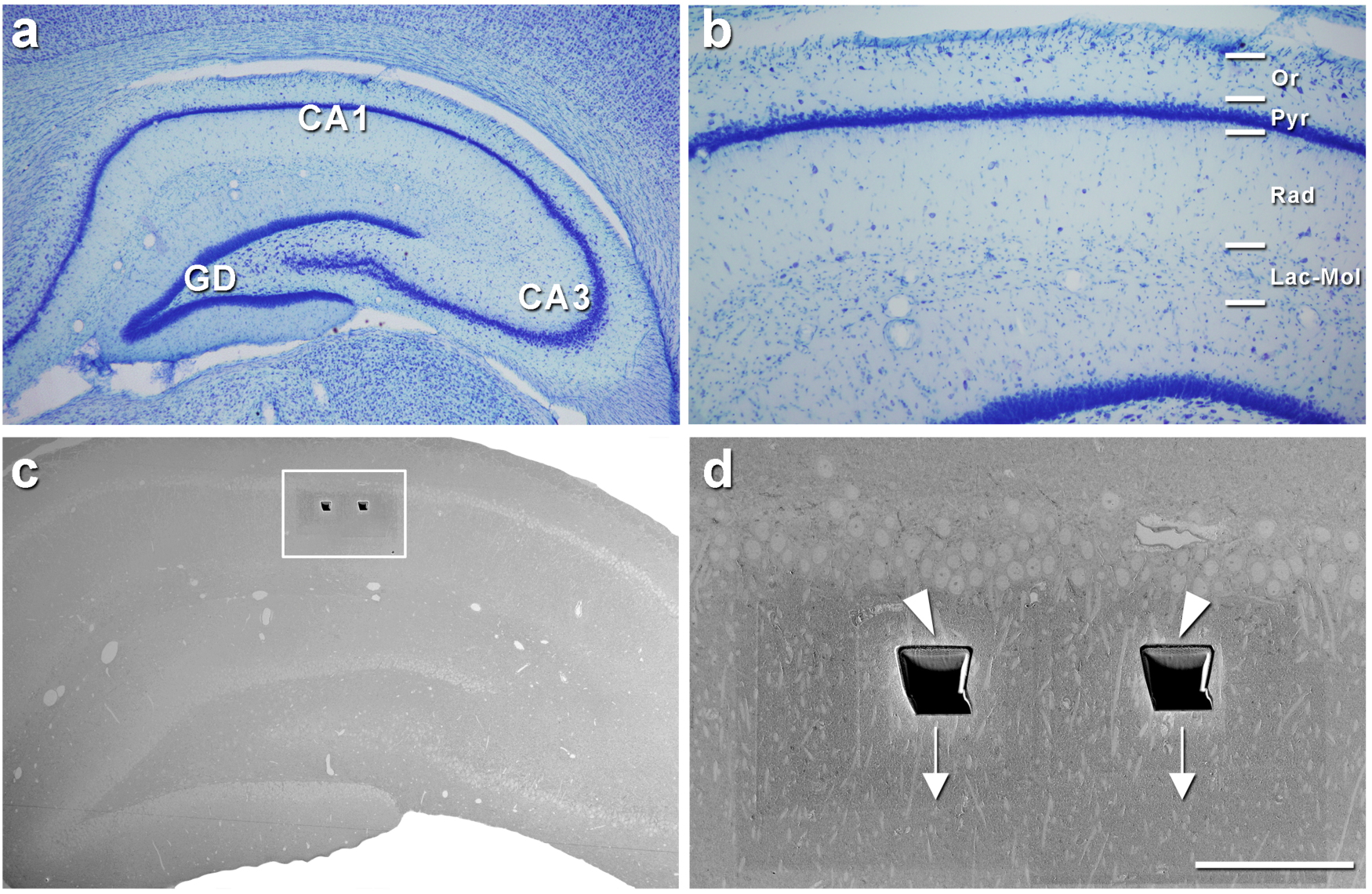
FIB/SEM imaging in the *stratum radiatum* of the rat CA1. **a**, Low-magnification photomicrograph of a coronal Nissl-stained brain section containing the hippocampus (CA1 and CA3: *Cornu ammonis* 1 and 3, GD: dentate gyrus). **b**, Higher magnification of the CA1 regions showing the layers (Or: *stratum oriens*, Pyr: *stratum pyramidale*, Rad: *stratum radiatum*, Lac-Mol: *stratum Lacunosum-Moleculare*). **c**, Image taken with the SEM from the surface of the block containing the hippocampus to be analzyed. Rectangle surrounds the region shown in **d. d**, Image taken with the SEM showing —at higher magnification— the *stratum pyramidale* and the trenches (arrowheads) opened with the FIB in the *stratum radiatum* to acquire the FIB/SEM serial images. Arrows indicate the direction of the FIB/SEM imaging. Scale bar in **d** corresponds to: 1 mm in **a**, 412 µm in **b**, 625 µm in **c**, 100 µm in **d**.

**Figure 2.**
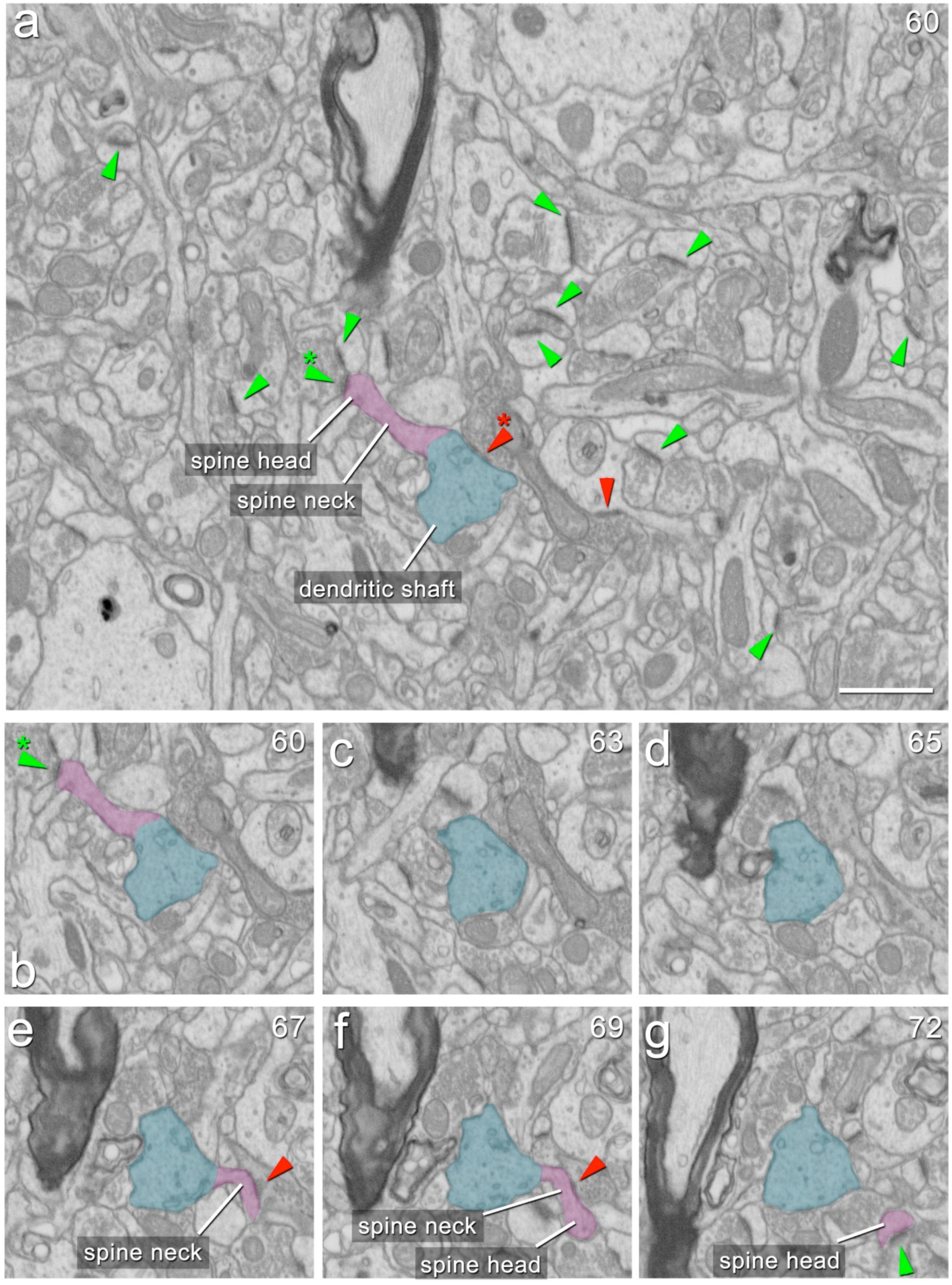
Differentiation of asymmetric and symmetric synapses and postsynaptic targets in the FIB/SEM stack of images. **a**, Example of an FIB/SEM image from a stack of images. Some asymmetric and symmetric synapses have been pointed out (green and red arrowheads, respectively). In addition, a spiny dendritc shaft (blue) with a dendritic spine (purple) emerging from the shaft is shown. The neck and the head of the spine have been indicated. A symmetric synapse on the dendritic shaft is pointed out with a red arrowhead. An axospinous asymmetric synapse (green arrowhead) is established on the head of the spine. **b**, A crop from **a** showing the same dendritic shaft and spine. **c**−**g**, Serial FIB/SEM images from the same cropped area in **b** to illustrate another dendritic spine emerging from the same shaft, by following up the stack of images. Note that in some images only the spine neck is observed (**e**), while in others it is possible to see only the spine head with no connection between it and the dendritic shaft (**g**). A symmetric synapse on the spine neck (red arrowhead) is observed in **e** and **f**. An axospinous asymmetric synapse (green arrowhead) is observed in **g**. The number of the section is indicated in the top right-hand corner of each section. Scale bar in **a** corresponds to: 1.3 µm in **a**, 1.4 µm in **b**−**g**.

**Figure 3.**
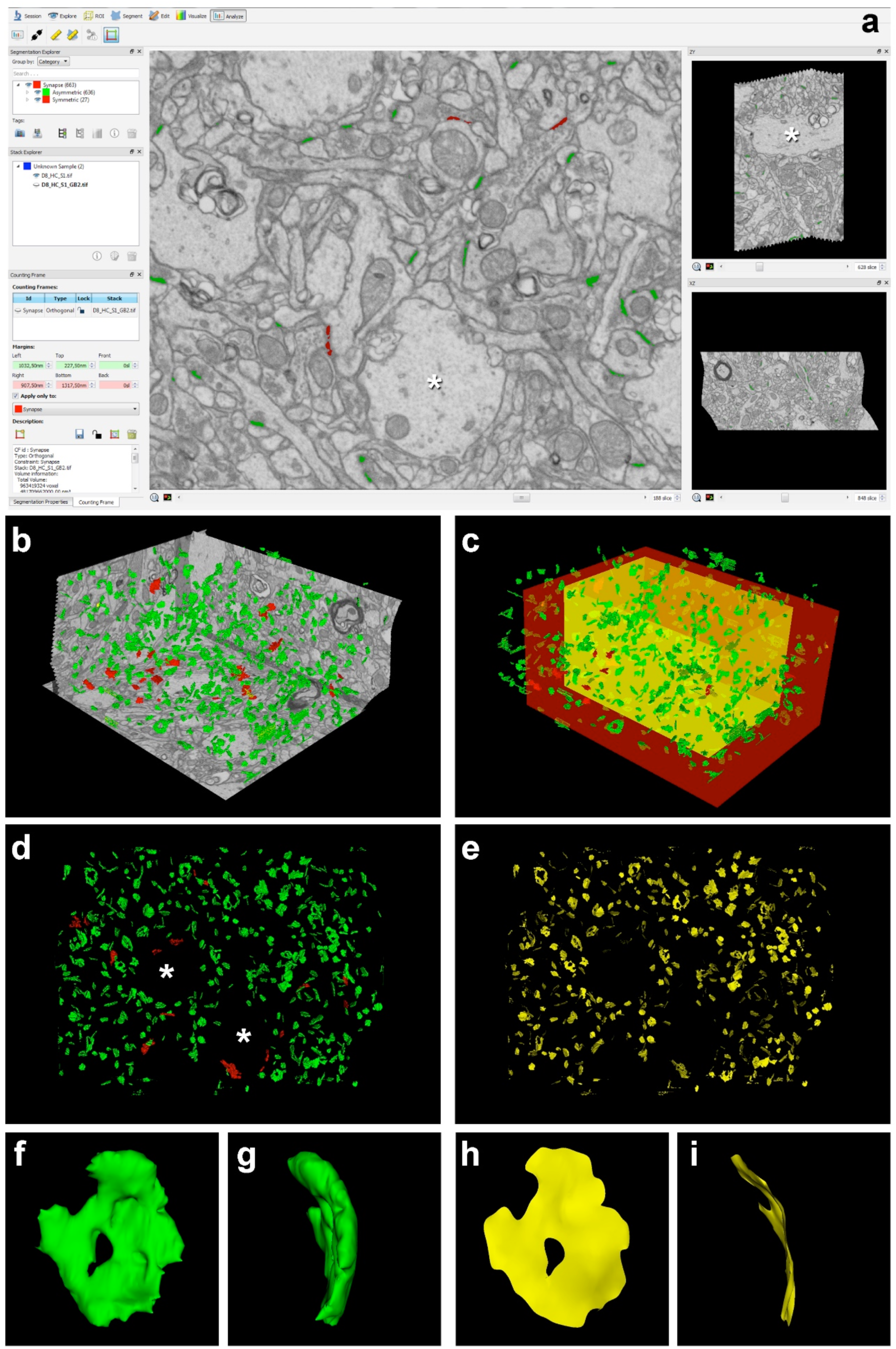
Analysis of the FIB/SEM stack of images with EspINA software. **a**, Screenshot from the EspINA software interface, which allows visualization of the image stacks and permits the identification and 3D reconstruction of all synapses in all spatial plans (XY, XZ and YZ). Asterisks point out large apical dendrites observed in the FIB/SEM stack of images: in the center, a coronally cut apical dendrite is observed and, on the right, the same dendrite in a lateral view, showing its elongated shape. **b**, 3D view of the three orthogonal planes and the 3D reconstruction of segmented synapses. **c**, Synapses are represented together with the counting frame (CF). Only synapses inside the inclusion volume of the CF are analyzed to ensure that all of them are complete (the exclusion area is shown in red; the inclusion area in yellow). **d**, Frontal 3D view of reconstructed synapses. Asterisks point out two spaces free of synapses due to the presence of large apical dendrites in the stack of images. **e**, Same frontal 3D view as in **d** but showing the SAS extracted from each reconstructed synapse. **f, g**, Two different 3D views of the same reconstructed asymmetric synapse. **h, i**, Same 3D views as in **f, g**, respectively, but showing the SAS extracted from the reconstructed synapse. Note the presence of a perforation in this synapse. Asymmetric synapses are colored in green and symmetric synapses in red.

### Synaptic density and ratio of asymmetric (AS) and symmetric (SS) synapses

After discarding incomplete and excluded synapses via the counting frame, a total of 19,317 synapses were analyzed, of which 10,990 synapses were from the saline group (10,561 AS and 429 SS; total tissue volume analyzed: 4,239 µm^3^, **Table 1**) and 9,069 from the cocaine-SA group (8,756 AS and 313 SS; total tissue volume analyzed: 3,774 µm^3^, **Table 1**). The number of synapses per unit volume (synaptic density) was calculated by dividing the total number of synapses by the volume of the counting frame.

**Table 1.** Data from the ultrastructural analysis of synapses in the neuropil of the *stratum radiatum* of the rat CA1 field in the saline and the cocaine-SA group. AS: asymmetric synapses; CA: *cornu Ammonis*; CF: counting frame; No.: number; SA: self-administration; SAS: synaptic apposition surface; SEM: standard error of the mean; SS: symmetric synapses. Data in parentheses are not corrected with the shrinkage factor.

|  | Saline | Cocaine-SA |
| --- | --- | --- |
| **No. synapses** | 10990 | 9069 |
| **No. AS** | 10561 | 8756 |
| **No. SS** | 429 | 313 |
| **% AS** | 96.25 | 96.54 |
| **% SS** | 3.75 | 3.46 |
| **CF volume ($\mu\text{m}^3$)** | 4239<br>(4427) | 3774<br>(3941) |
| **No. synapses / $\mu\text{m}^3$**<br>**(mean ± SEM)** | 2.52±0.13<br>(2.63±0.14) | 2.29±0.13<br>(2.39±0.14) |
| **No. AS / $\mu\text{m}^3$**<br>**(mean ± SEM)** | 2.42±0.13<br>(2.53±0.13) | 2.21±0.13<br>(2.31±0.14) |
| **No. SS / $\mu\text{m}^3$**<br>**(mean ± SEM)** | 0.094±0.008<br>(0.099±0.009) | 0.078±0.003<br>(0.081±0.003) |
| **Area of AS SAS**<br>**($\text{nm}^2$; mean ± SEM)** | 42022±2310<br>(40206±2422) | 48916±3249<br>(47522±3156) |
| **Area of SS SAS**<br>**($\text{nm}^2$; mean ± SEM)** | 67972±6025<br>(66256±5825) | 90738±9799<br>(88153±9520) |
| **Perimeter of AS SAS**<br>**(nm; mean ± SEM)** | 996 ± 32<br>(981 ± 32) | 1077 ± 34<br>(1062 ± 34) |
| **Perimeter of SS SAS**<br>**(nm; mean ± SEM)** | 1507 ± 81<br>(1485 ± 80) | 1916 ± 153<br>(1888 ± 151) |
| **Curvature of AS SAS**<br>**(mean ± SEM)** | 0.0037 ± 0.0003 | 0.0034 ± 0.0002 |
| **Curvature of SS SAS**<br>**(mean ± SEM)** | 0.0028 ± 0.0004 | 0.0031 ± 0.0007 |
| **Volume fraction of**<br>**mitochondria (%; mean ± SEM)** | 7.31 ± 0.20 | 7.49 ± 0.23 |

The synaptic density (including both AS and SS) in the saline group was 2.52 synapses/µm^3^, comprising 2.42 and 0.094 synapses/µm^3^ for AS and SS, respectively; that is, the ratio of AS to SS was 96.25:3.75 (**Figure 4a,b, Table 1**).

**Figure 4.**
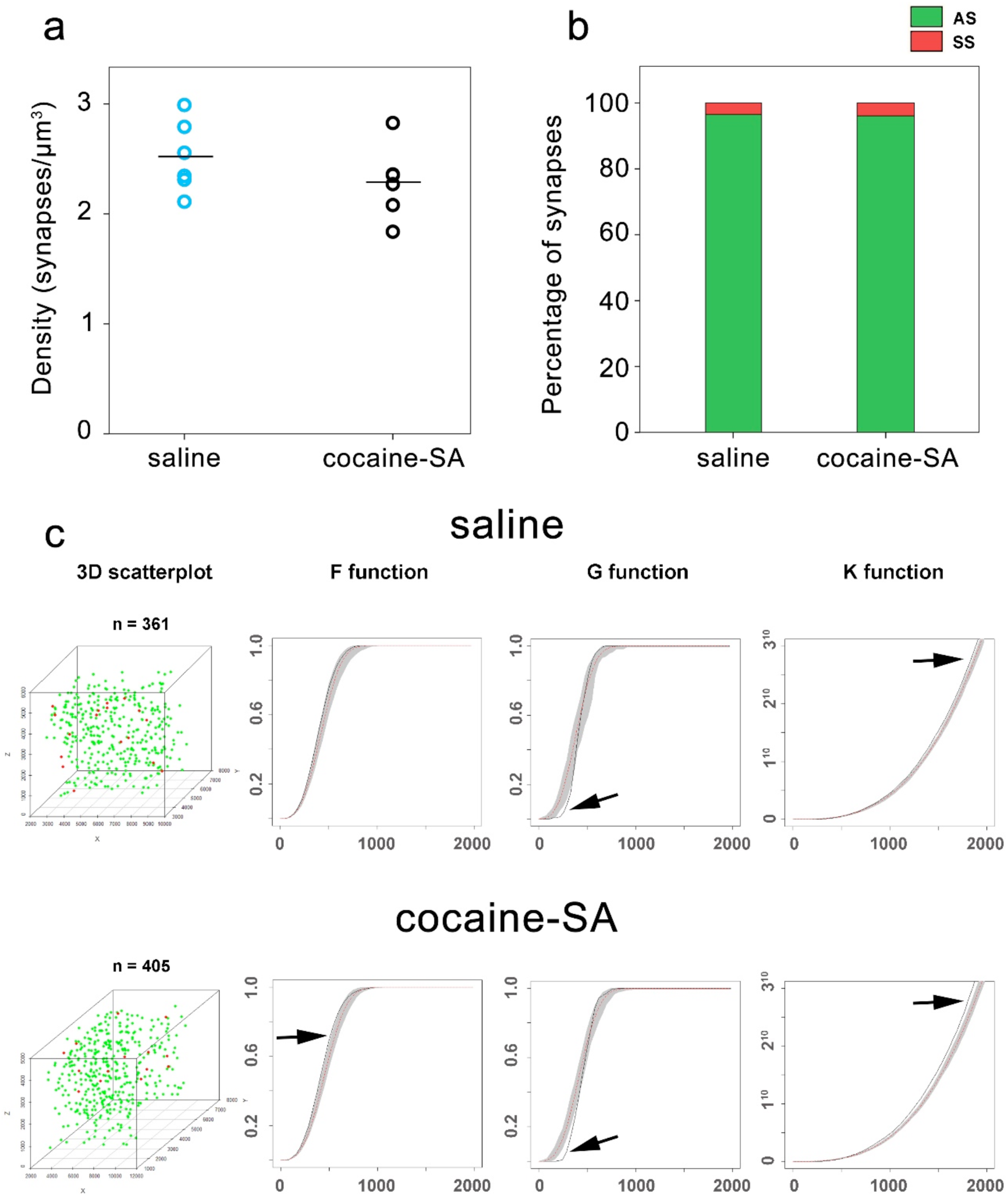
Synaptic density, proportions and spatial distribution in the *stratum radiatum* of the rat CA1 in the saline and the cocaine-SA groups. **a**, Graph showing the mean synaptic density (synapses / µm^3^) of all synapses. Each circle corresponds to the mean synaptic density value in each animal (n=6 in the saline group and n=6 in the cocaine-SA group). **b**, Graph showing the percentage of asymmetric and symmetric synapses in the saline and the cocaine-SA group. **c**, Example of the synaptic spatial distribution analysis in two samples — one from the saline group and the other from the cocaine-SA group (upper and bottom rows, respectively). For each synapse, we recorded the spatial positions of the center of gravity or centroid of the synaptic junction, as represented in the 3D scatterplots of the leftmost column (with the number of synapses analyzed shown above each of the two scatterplots). Three spatial statistical functions (F, G and K) were calculated for each group of synapses in each sample. In each graph, the function corresponding to the actual sample is represented by a black line. The theoretical homogeneous Poisson distribution or complete spatial randomness (CSR) is represented as a red discontinuous trace, and the grey envelope is generated by 99 simulations of the CSR model. The spatial distribution of all synapses was nearly random, since all three functions lay within the simulated envelopes or deviated only slightly (arrows). Deviations in G functions occur because synaptic junctions cannot overlap, and thus the minimum intersynaptic distances (measured between their centroids) must be limited by the sizes of synaptic junctions themselves. Deviations in F and K functions can be due to the presence of large apical dendrites in this layer of CA1 that leave spaces free of synapses (see **Figure 3a,d**). No differences were observed in the synaptic density (t_10_=1.208, p=0.255), the proportion of asymmetric or symmetric synapses (χ^2^=2.853, p=0.091), or the spatial distribution of synpases after cocaine-SA. AS: asymmetric synapses; SA: self-administration; SS: symmetric synapses.

No differences were found after cocaine-SA regarding synaptic density (including both AS and SS) (2.29 synapses/µm^3^ in cocaine-SA group; t_10_=1.208, p=0.255; **Figure 4a, Table 1**) or when analyzing the densities of AS and SS separately (AS: 2.21 synapses/µm^3^ in cocaine-SA group, t_10_=1.145, p=0.279; and SS: 0.078 synapses/µm^3^ in cocaine-SA group, t_10_=1.870, p=0.091; **Table 1**). The ratio of AS to SS was also unaltered after cocaine-SA (96.54:3.46 in cocaine-SA group, χ^2^=2.853, p=0.091; **Figure 4b; Table 1**).

### Synaptic spatial distribution

An analysis of the spatial distribution of synapses was performed for all synapses (AS and SS together). Our results indicate that the spatial organization of synapses in the neuropil of the *stratum radiatum* is nearly random, since only slight deviations from the CSR model were found (**Figure 4c**). No significant differences were observed in the spatial distribution of synapses between the saline and the cocaine-SA group (**Figure 4c**).

### Distribution of postsynaptic targets

Two main postsynaptic targets were considered (**Figure 2**): dendritic spines (axospinous synapses) and dendritic shafts (axodendritic synapses). In the case of axospinous synapses, the exact location of the synaptic contact was determined (i.e., the head or neck of the dendritic spine). For axodendritic synapses, dendritic shafts were further classified as spiny (when dendritic spines could be observed emerging from the shaft) or aspiny. Only synapses whose postsynaptic target was clearly identifiable after navigation through the stack of images (saline group: n=5,029; AS=4,818, SS=211; cocaine-SA group: n=4,708; AS=4,535, SS=173) were considered for analysis (**Supplementary table 1**).

#### Saline group

##### Total synaptic population

Most synapses (AS+SS) were established on dendritic spines —especially on the head— (n=4,775; 94.94 %) rather than on dendritic shafts (n=254; 5.06 %) (χ^2^=4,064.315, p=0.000). Synapses (AS+SS) on spiny shafts were more abundant than synapses on aspiny shafts (4.11 and 0.95 %, respectively, χ^2^=8,599.176, p=0.000) (**Figure 5a**).

**Figure 5.**
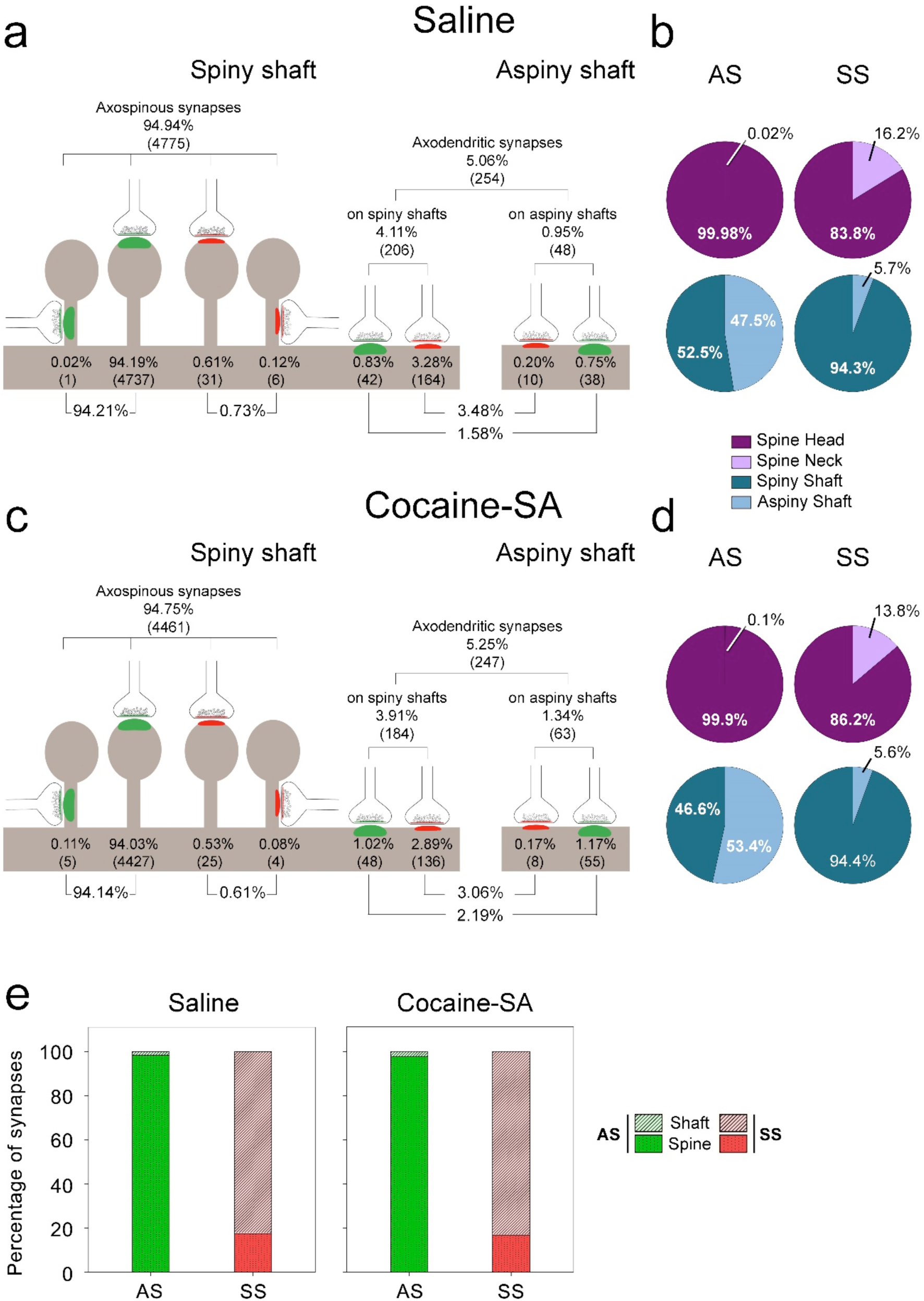
Distribution of synapses according to their postsynaptic targets in the *stratum radiatum* of the rat CA1 in the saline and cocaine-SA groups. **a, c**, Percentages of axospinous (both on the head and neck of dendritic spines) and axodendritic (both on spiny and aspiny shafts) asymmetric synapses (AS; green) and symmetric synapses (SS; red) in the saline and the cocaine-SA group, respectively. The numbers of each synaptic type are shown in brackets. **b, d**, Pie charts illustrating the proportions of AS and SS according to their location as axospinous synapses (i.e., on the head or neck of the spine) or axodendritic synapses (i.e., spiny or aspiny shafts) in the saline and the cocaine-SA group, respectively. **e**, Graph showing the percentage of axospinous and axodendritic synapses within the AS and SS populations in the saline and the cocaine-SA groups. AS: asymmetric synapses; SA: self-administration; SS: symmetric synapses.

As a whole, axospinous AS were clearly the most abundant type of synapse (n=4,738; 94.21 %), followed by axodendritic SS (n=174; 3.48 %), axodendritic AS (n=80; 1.58 %) and axospinous SS (n=37; 0.73 %) (**Figure 5a**).

##### Postsynaptic preference of AS and SS

Most AS were established on dendritic spines (n=4,738, 98.34 %; **Figure 5e, Supplementary table 1**) and almost exclusively on the head of the spines (99.98%; **Figure 5b**). The remaining AS were established on dendritic shafts (n=80, 1.66% — 52.5% of which were on spiny dendritic shafts and 47.5% on aspiny shafts; **Figure 5b,e, Supplementary table 1**). By contrast, most SS were axodendritic (n=174, 82.47%; **Figure 5e, Supplementary table 1**) and had a clear preference for spiny shafts versus aspiny shafts (94.3% on spiny dendritic shafts and 5.7% on aspiny shafts; **Figure 5b**). A lower percentage of SS were established on dendritic spines (n=37, 17.53 %; **Figure 5e, Supplementary table 1**) and they were found more commonly on the head of the spines than on spine necks (83.8% of AS were on spine heads and 16.2% were on spines necks; **Figure 5b**).

The preference of AS and SS for dendritic spines and dendritic shafts, respectively, was statistically tested. We found a consistent association (χ^2^= 2,752.261, p=0.000) for AS and dendritic spines (receiving 99.2% AS and 0.8% SS), and for SS and dendritic shafts (receiving 31.5% AS and 68.5% SS). Regarding dendritic spines, we found that AS had a significant preference for spine heads: 99.3% of AS and only 0.7% of SS (χ^2^ = 2,873.067, p=0.000) were on spine heads, while SS had a significant preference for spine necks (χ^2^=115.886, p=0.000) (with 85.7% SS established on spine necks versus 14.3% in the case of AS). Regarding dendritic shafts, SS presented a significant preference both for spiny dendritic shafts (receiving 20.4% AS and 79.6% SS; χ^2^ = 3,039.292, p=0.000) and aspiny dendritic shafts (establishing 79.2% AS and 20.8% SS; χ^2^=33.374, p=0.000) (**Supplementary tables 2, 3**).

See Supplementary Results for further information about multiple synapses per spine head.

#### Cocaine-SA group

The distribution of postsynaptic targets after cocaine-SA was similar to the saline group with only slight differences as described below (**Figure 5, Supplementary table 1**).

We observed slight differences in the distribution of postsynaptic targets after cocaine-SA in the case of AS (χ^2^=7.916, p=0.043) but not for SS (χ^2^=0.119, p=0.989 for SS). Specifically, following cocaine-SA, we found a slightly lower proportion of AS on spine heads (dropping from 98.32% to 97.62% — a decrease of 0.7%, χ^2^=5.786, p=0.016) and a slightly higher proportion of AS on aspiny shafts (rising from 0.79% to 1.21%: an increase of 53%, χ^2^=4.268, p=0.039) (**Supplementary table 1**).

### Synaptic shape

EspINA software was used to measure the synaptic apposition surface (SAS) in 3D, and classify them into 4 morphological categories: macular, horseshoe-shaped, perforated and fragmented (**Figure 6a**). We determined the synaptic shape of 5,117 synapses (4,902 AS and 215 SS) in the saline group and 4,767 synapses (4,589 AS and 178 SS) in the cocaine-SA group (**Supplementary table 4**).

**Figure 6.**
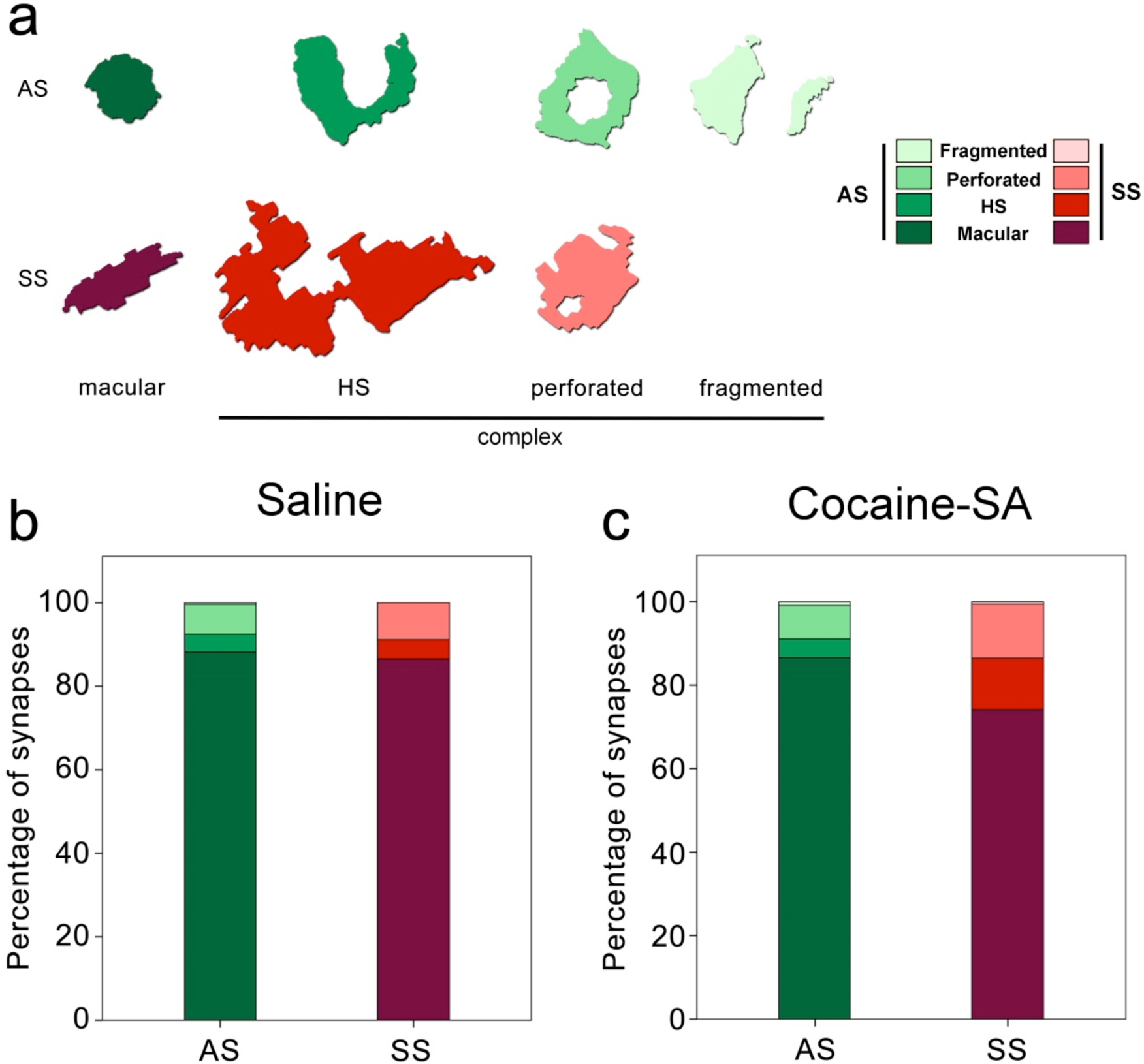
Proportions of synaptic shapes in the *stratum radiatum* of the rat CA1 in the saline and cocaine-SA groups. **a**, Examples of the different types of synapses based on the shape of the synaptic junction: macular, horseshoe-shaped (HS), perforated and fragmented. We also refer to non-macular synapses as synapses with complex shapes. The upper and lower rows show examples of shapes of AS and SS, respectively. **b, c**, Percentages of the different types of synaptic shapes within the population of AS and SS in the saline (**b**) and the cocaine-SA (**c**) group. AS: asymmetric synapses; HS: horseshoe-shaped synapses; SA: self-administration; SS: symmetric synapses.

#### Saline group

The four synaptic shapes were not equally present within AS and SS (χ^2^=10,497.767, p=0.000 and χ^2^=274.167, p=0.000, respectively). The vast majority of both AS and SS had a macular shape (88.23% and 86.51%, respectively), followed by perforated synapses (7.16% and 8.84%, respectively), horseshoe-shaped synapses (4.24% and 4.65%, respectively) and fragmented synapses (0.37% and 0%, respectively, **Figure 6b; Supplementary table 4**). When grouping horseshoe-shaped, perforated and fragmented synapses as synapses with complex shapes, we found that macular synapses were more frequently found than those with more complex shapes, both for AS (macular: 88.23% and complex: 11.77%; χ^2^=2,865.668, p=0.000) and SS (macular: 86.51% and complex: 13.49%; χ^2^=114.647, p=0.000).

We did not observe an association between the type of synapse (AS or SS) and the synaptic shape (χ^2^ = 1.740, p=0.628).

We did not observe an association between the type of postsynaptic target and the synaptic shape for either AS (χ^2^=3.149, p=0.338) or SS (χ^2^=2.461, p=0.277) (**Supplementary table 5**). Thus, the synaptic shape distribution was the same for axospinous and axodendritic synapses — both for AS and SS.

#### Cocaine-SA group

The proportion of synaptic shapes after cocaine-SA was similar to the saline group and only slight differences were found, which we describe below (**Figure 6, Supplementary table 4**).

We observed that the distribution of synaptic shapes was different after cocaine-SA for AS (χ^2^=13.916, p=0.003) and SS (χ^2^=11.531, p=0.005). Specifically, for AS, we found that after cocaine-SA, macular AS were less frequently found (declining from 88.23 to 86.60 — a decrease of 1.8%, χ^2^=5.741, p=0.017) and fragmented AS were more frequently present (rising from 0.37 to 0.89 — an increase of 140%, χ^2^=10.625, p=0.001). For SS, we found that after cocaine-SA, macular SS were less frequently found (they dropped from 86.51 to 74.16 — a decrease of 14%, χ^2^=9.625, p=0.002) and horseshoe-shaped SS were more frequently present (rising from 4.65 to 12.36 — an increase of 166%, χ^2^=7.736, p=0.005) (**Figure 6, Supplementary table 4**). Regarding the postsynaptic targets, we observed that these differences were found within axospinous synapses in the case of AS (χ^2^=14.812, p=0.002), and within axodendritic synapses in the case of SS (χ^2^=9.210, p=0.027) (**Supplementary table 5**).

### The synaptic apposition surface (SAS)

Morphological features of the SAS were obtained with EspINA software for both AS and SS. We analyzed the SAS of 10,990 synapses in the saline group (AS=10,561; SS=429) and 9,069 synapses in the cocaine-SA group (AS=8,756; SS=313).

### Synaptic size: SAS area and perimeter

#### Saline group

SAS areas ranged from 1990 to 318,912 nm^2^ for AS, and from 3,634 to 620,661 nm^2^ for SS. The mean SAS area of AS (42,022 nm^2^) was significantly smaller than the mean SAS area of SS (67,972 nm^2^) (t_10_=-4.022, p=0.002) (**Figure 7a, Table 1**).

**Figure 7.**
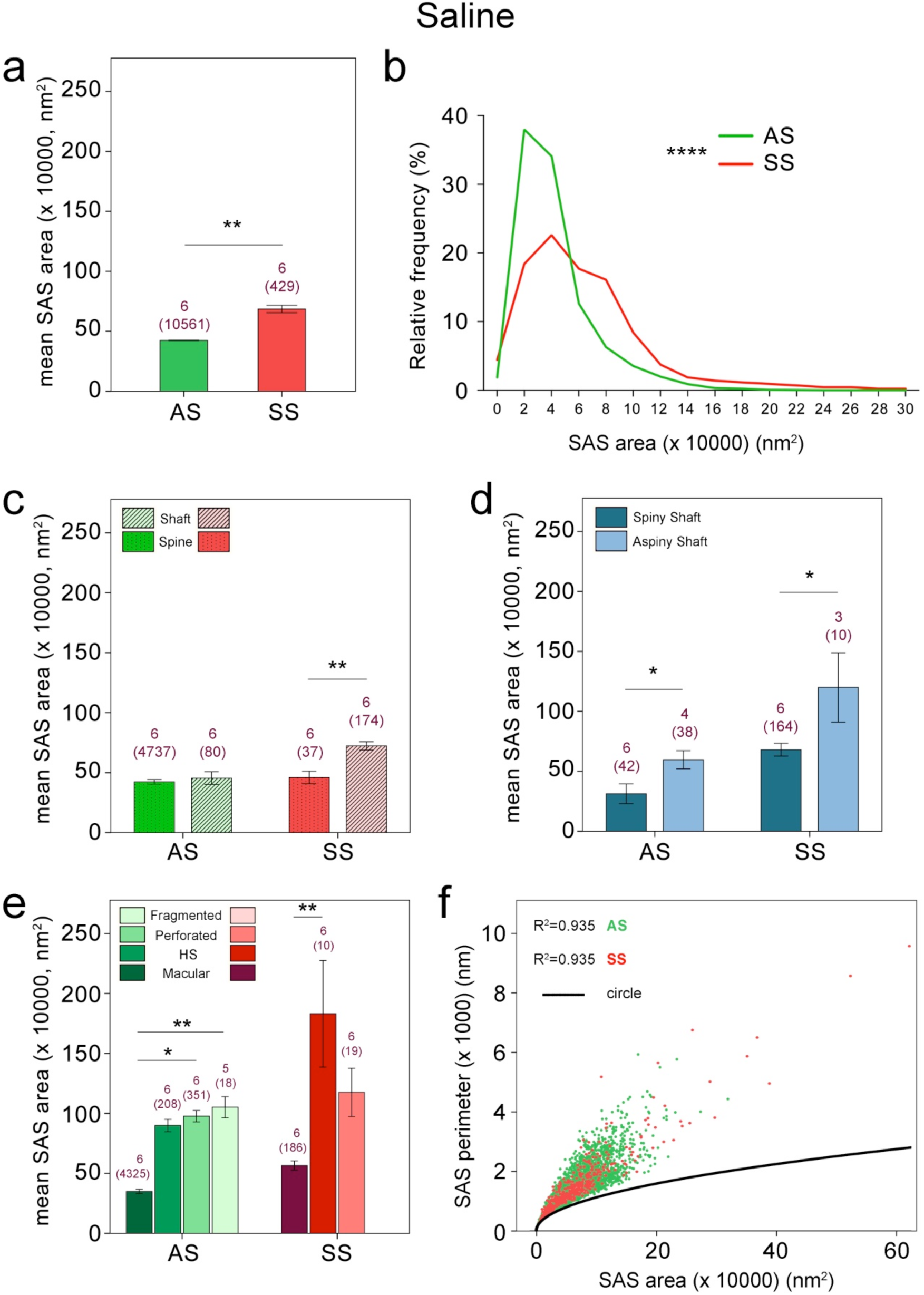
Synaptic apposition surface (SAS) area measurements in the *stratum radiatum* of the rat CA1 in the saline group. **a**, Mean SAS area of asymmetric synapses (AS; green) and symmetric synapses (SS; red) are represented (mean±SEM). **b**, Frequency distribution of SAS areas for both AS (green line) and SS (red line). **c**, Mean SAS area of axospinous and axodendritic synapses are also shown for AS and SS (mean±SEM). **d**, Mean SAS area of axodendritic synapses on spiny and aspiny shafts are also shown for AS and SS (mean±SEM). **e**, Mean SAS area related to the different synaptic shapes are plotted for both AS and SS (mean±SEM). **f**, Scatter plot showing the relationship between SAS areas and perimeters. AS are represented as green dots and SS as red dots. The black trace indicates the perimeter/area relation of a circle, as a reference. There is a strong correlation between SAS area and perimeter for AS (R^2^=0.935) and SS (R^2^=0.935). The number of animals and the number of synapses analyzed (in brackets) is indicated. AS: asymmetric synapses; HS: horseshoe-shaped synapses; SEM: standard error of the mean; SS: symmetric synapses. *p<0.05, **p<0.01, ***p<0.001, see text for further information about statistical comparisons.

Mean perimeters were larger for SS than for AS (996 nm and 1,507 nm, respectively, t_10_=-5.836, p=0.000) (**Table 1**). There was a strong correlation between SAS area and perimeter for (i) all synapses (R^2^=0.936, p=0.000), (ii) AS (R^2^=0.935, p=0.000) and (iii) SS (R^2^=0.935, p=0.000) (**Figure 7f**). Therefore, the larger the SAS area of a synapse, the more tortuous its perimeter — the SAS perimeter tends to grow faster than the perimeter of a circle (**Figure 7f**).

To further characterize the size distribution of SAS, we plotted the frequency histograms of SAS areas and perimeters. For AS and SS, these frequency histograms showed a unimodal and continuous distribution with a positive skewness. The SAS area and perimeter distribution of AS statistically differed from SS (KS=5.927, p=0.000 for area; KS=7.768, p=0.000 for perimeter) and a longer right tail was observed for SS (**Figure 7b**).

Additionally, we studied the synaptic size according to the postsynaptic targets (**Figure 7c,d; Supplementary table 6**). For AS, axospinous AS were the same size as axodendritic AS, while —for SS— axospinous SS were smaller than axodendritic SS. When focusing on the type of dendritic shaft, we observed that synapses on aspiny shafts were larger than synapses on spiny shafts, both for AS and for SS. Finally, we analyzed the differences in the synaptic size according to the shape of the synaptic junctions (**Figure 7e; Supplementary table 7**). Macular AS were smaller than the rest of the synaptic shapes within AS, while macular SS were smaller than horseshoe-shaped SS. See Supplementary results for information about statistical significances.

#### Cocaine-SA group

No differences were found in the mean SAS area or perimeter of AS and SS after cocaine-SA (AS: t_10_=-1.729, p=0.114 for area and t_10_=-1.736, p=0.113 for perimeter; SS: t_10_=-1.979, p=0.076 for area), with the exception of SS with larger perimeters (saline: 1507 µm; cocaine-SA: 1916 µm, t_10_=-2.357, p=0.040) (**Table 1**). In addition, we did not find significant differences after cocaine-SA in the mean SAS area or perimeter of AS and SS classified according to the postsynaptic targets or the synaptic shapes (**Supplementary tables 6, 7**).

However, when analyzing the SAS area and perimeter distribution, the cocaine-SA group showed a higher number of synapses with larger SAS area and perimeter than the saline group, both for AS (KS=6.550, p=0.000 for area; KS=4.885, p=0.000 for perimeter) and for SS (KS=4.430, p=0.000 for area; KS=2.790, p=0.000 for perimeter) (**Figure 8a,b**). When focusing on the postsynaptic targets, a higher number of synapses with larger SAS area and perimeter was observed for axospinous AS after cocaine-SA (KS=3.715, p=0.000 for area; KS=2.636, p=0.000 for perimeter), while these size differences were less evident —or even non-existent— for axodendritic AS (KS=2.025, p=0.001 for area; KS=1.323, p=0.060 for perimeter) (**Figure 8c,e**). No size differences were observed for axospinous SS (KS=0.800, p=0.543 for area; KS=0.782, p=0.574), but there was a slight tendency for a higher number of synapses with larger SAS area and perimeter for axodendritic SS after cocaine-SA (KS=1.334, p=0.057 for area; KS=1.359, p=0.050 for perimeter) (**Figure 8d,f**). Regarding synaptic shape, a higher number of synapses with larger SAS area and perimeter was observed within the population of macular AS after cocaine-SA (KS=3.835, p=0.000 for area; KS=2.583, p=0.000 for perimeter), while these size differences were less evident or even non-existent for horseshoe-shaped AS (KS=1.770, p=0.004 for area; KS=1.313, p=0.064 for perimeter). No changes were observed for perforated and fragmented AS (Perforated AS: KS=1.227, p=0.098 for area and KS=0.665, p=0.769 for perimeter; Fragmented AS: KS=0.585, p=0.884 for area and KS=0.867, p=0.439 for perimeter) (**Figure 9a,c,e,g**). For SS, only the population of perforated SS showed a slight tendency for a higher number of larger synapses after cocaine-SA (Macular SS: KS=0.852, p=0.462 for area and KS=0.936, p=0.346 for perimeter; Horseshoe-shaped SS: KS=0.731, p=0.659 for area and KS=0.818, p=0.515 for perimeter; Perforated SS: KS=1.484, p=0.024 for area and KS=1.144, p=0.146 for perimeter) (**Figure 9b,d,f**). Considering the figures for significance (p>0.001 for AS), together with both the results for SAS area and perimeter distribution, we interpret that the higher proportion of larger synapses after cocaine-SA was especially evident for axospinous and macular AS, while —in the case of SS— the greater proportion of larger synapses was not so clearly associated with any specific synaptic type.

**Figure 8.**
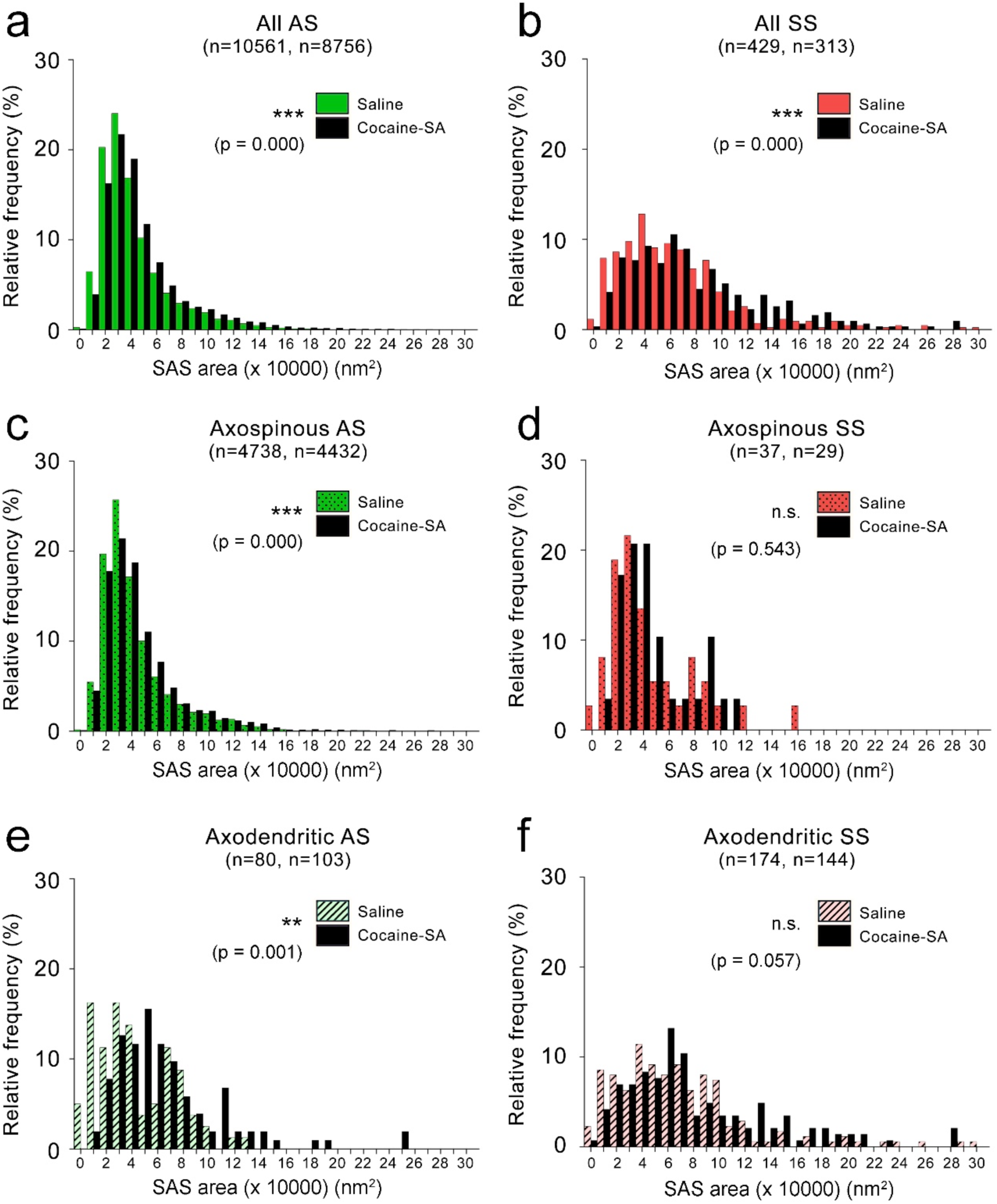
Frequency distribution of the synaptic apposition surface (SAS) areas for all synapses classified according to their postsynaptic targets in the saline and cocaine-SA groups. **a, b**, Frequency distribution of SAS areas for both AS (**a**) and SS (**b**). **c, d**, Frequency distribution of SAS areas for both axospinous AS (**c**) and axospinous SS (**d**). **e, f**, Frequency distribution of SAS areas for both axodendritic AS (**e**) and axodendritic SS (**f**). The number of synapses in the saline and the cocaine-SA group, respectively, is indicated in brackets. AS: asymmetric synapses; SA: self-administration; SS: symmetric synapses. **p<0.01, ***p<0.001, see text for further information about statistical comparisons.

**Figure 9.**
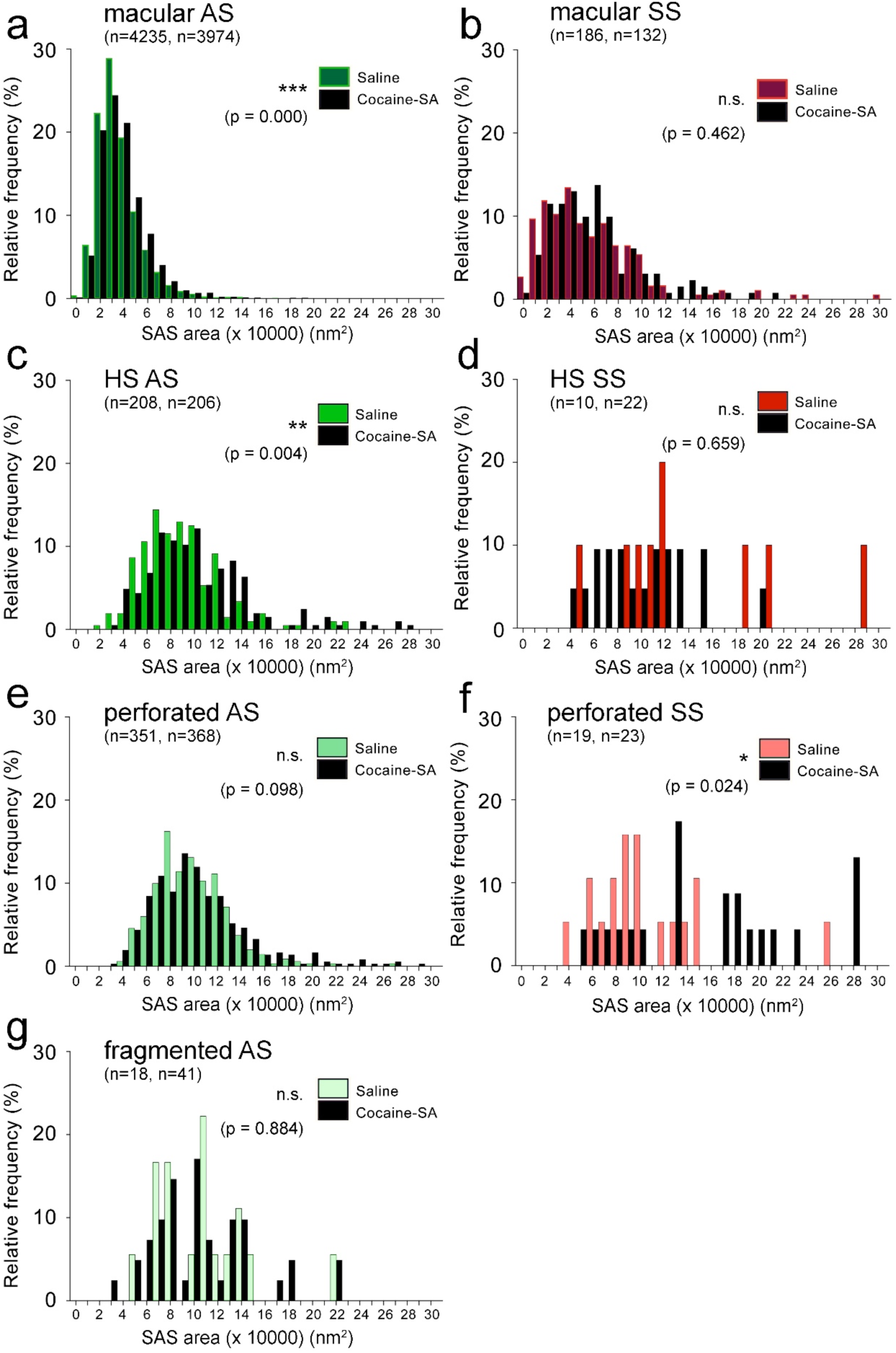
Frequency distribution of the synaptic apposition surface (SAS) areas for synapses classified according to their synaptic shape in the saline and cocaine-SA groups. **a, b**, Frequency distribution of SAS areas for both macular AS (**a**) and macular SS (**b**). **c, d**, Frequency distribution of SAS areas for both horseshoe-shaped AS (**c**) and horseshoe-shaped SS (**d**). **e, f**, Frequency distribution of SAS areas for both perforated AS (**e**) and perforated SS (**f**). **g**, Frequency distribution of SAS areas for fragmented AS. The number of synapses in the saline and the cocaine-SA group, respectively, is indicated in brackets. AS: asymmetric synapses; HS: horseshoe-shaped synapses; SA: self-administration; SS: symmetric synapses. *p<0.05, **p<0.01, ***p<0.001, see text for further information about statistical comparisons.

See Supplementary Results for information about the curvature of the synapses and the volume fraction of neuropil occupied by mitochondria.

## DISCUSSION

We can draw the following two main conclusions from our work:

1. In the control rats, most synapses are AS and randomly distributed. The most common synaptic type is an AS with a macular shape on a spine head. This type of synapse is also smaller than the rest of the synaptic types.
2. In the cocaine-SA rats, there are no changes in the synaptic density, the ratio of AS and SS, or in the synaptic spatial distribution. However, we observed some differences in the distribution of postsynaptic targets and synaptic shapes as well as in the synaptic size. A higher proportion of larger AS (mainly, axospinous and macular) and SS were observed. This increase in the synaptic size was accompanied by the presence of a higher proportion of synapses with more complex shapes (that have larger sizes than macular ones).

A summary of the results is shown in **Supplementary figure 2** and **Supplementary table 8**.

### Synaptic organization of the *stratum radiatum* of the control rat CA1 (saline group)

#### Synaptic density, ratio of AS and SS, and spatial distribution

##### Synaptic density

We found a synaptic density of 2.52 synapses / µm^3^. Using FIB/SEM, we have previously shown that the synaptic density in the *stratum radiatum* of CA1 of the mouse is 2.36 synapses / µm^3^ ^(29)^, whereas in the *stratum radiatum* of CA1 of the human hippocampus, it is 0.67 synapses / µm^3^ ^(30)^. Thus, the synaptic density of the rat and mouse is remarkably similar but there are huge differences in synaptic density between humans and rodents. These differences —together with differences in other anatomical, genetic, molecular and physiological features (e.g.,^31,32,33^)— further support the notion that remarkable differences exist between the human and rodent CA1. These differences clearly need to be taken into consideration when making interpretations in translational studies comparing one species to another.

##### Ratio of AS and SS

It has been consistently reported that the neuropil is characterized by a much higher number of excitatory contacts compared to inhibitory synapses in different brain regions and species^34,35,36,37,38,39,24^. In the present study, we found that the AS:SS ratio was 96.25:3.75. Preliminary results from our laboratory, using the FIB/SEM methodology, have shown that the AS:SS ratios are 98:2 and 96:4 in the *stratum radiatum* of the mouse and human CA1, respectively^29,30^. That is, the ratio of AS and SS in the CA1 field is similar in humans and rodents.

##### Spatial distribution

We found that the spatial organization of synapses was nearly random, with slight deviations which could be explained by the presence of large apical dendrites in *stratum radiatum* that leave relatively large spaces free of synapses (see **Figure 3d**). Randomly distributed synapses have been described in the somatosensory cortex of normal rats and the frontal and transentorhinal cortices and hippocampus of the human brain^40,41,37,30^, suggesting that this synaptic characteristic is a widespread ‘rule’ of the cerebral cortex of different species. However, slight deviations from randomness —like those we observed in the rat hippocampus— were not observed in the human hippocampus.

#### Postsynaptic targets

The most abundant synaptic type was axospinous AS (94.21%), followed by axodendritic SS (3.48%) and axodendritic AS (1.58%). Axospinous SS were rarely observed (0.73%). Regarding postsynaptic preferences, most inhibitory synapses (82.47%) were on dendritic shafts, while most excitatory synapses (98.34%) were on spines (almost exclusively on the head of the spine). This preference of inhibitory and excitatory axons for dendritic shafts and dendritic spines, respectively, is also characteristic of other cortical regions and species, although variations in their percentages have been reported^34,42,43,39,35,36,24,38^.

Regarding axospinous AS, they are especially abundant (94.21%) when compared to other brain regions in both humans and other species^34,42,43,39,36,35,24,38,30^. For example, in the *stratum radiatum* of the human CA1 and the somatosensory cortex of the rat, axospinous AS represent around 75% of the total synaptic population^24,30^ and in layer II of the human transentorhinal cortex or the *stratum lacunosum-moleculare* of the human CA1, axospinous AS are even less numerous and account for around 55% of the total synaptic population^38,30^.

#### Synaptic shape, synaptic size and curvature

Most synapses presented a simple, macular shape (around 87% for both AS and SS), in agreement with previous reports in different brain areas and species^44,45,46,38,30^. In our study, non-macular synapses constituted around 13% of the total synapses. Similar results to ours have been found using FIB/SEM microscopy in the *stratum radiatum* of the human CA1 (13% non-macular AS and 18% non-macular SS^30^).

The shape and size of the synaptic junctions are strongly correlated with the release probability, synaptic strength, efficacy and plasticity^47,48,49,50^. In this regard, all three types of non-macular synapses (with more complex shapes) were larger than macular ones, as previously observed in different brain regions and species^46,38,30^. Although the functional significance of perforations is still unclear, perforated synapses are known to have more AMPA and NMDA receptors than macular synapses and are thought to constitute a relatively powerful population of synapses with more long-lasting memory-related functionality than their smaller, macular counterparts^47,48,51^.

Considering all synapses, inhibitory contacts were approximately 62% larger than excitatory ones — a similar trend to that observed in the rat somatosensory cortex (SS were 43% larger than AS in layers I−VI)^46^. However, this contrasted with our findings in layer II of the human transentorhinal cortex^37^ and the human CA1^30^; AS were 55% larger than SS in layer II of the transentorhinal cortex and 31% larger in the *stratum radiatum* of the CA1. Interestingly, AS are much larger (almost twice as big) in the *stratum radiatum* of the human CA1, while SS sizes in rats and humans are relatively similar^30^. For further discussion about similarities and differences regarding synaptic size and curvature between rat and human cortex, see Supplementary Discussion.

Therefore, the synaptic organization of the *stratum radiatum* of the rat CA1 shows remarkable differences compared to the same layer and region of the human hippocampus as well as compared to other brain regions and species.

### Effects of cocaine-SA on the synaptic organization of the *stratum radiatum* of the rat CA1

#### Synaptic density, ratio of AS and SS, and spatial distribution

No significant changes were found after cocaine-SA with regard to the synaptic density, the ratio of AS and SS, or the synaptic spatial distribution. However, previous light microscopy studies from our laboratory have shown that both passive administration of cocaine (for 15 days by intraperitoneal injection) and cocaine-SA for 3 weeks increase the number of dendritic spines in the *stratum radiatum* of the CA1 field of the Lewis rat hippocampus^19,13^, which a priori could be interpreted as an increase in the number of axospinous synapses in cocaine-SA rats. This discrepancy with our results might be explained by the possibility that newly appearing dendritic spines in cocaine-SA are non-synaptic, resulting in the increase in the spine number not being accompanied by a synaptic density increase. Thus, further studies are needed to determine if there is an increase in the proportion of non-synaptic dendritic spines in cocaine-SA animals.

Since no differences were found in the ratio of AS and SS after cocaine-SA, the balance between excitation and inhibition does not seem to be altered in the hippocampus following cocaine-SA. However, changes in synaptic size and activity can occur independently of synaptic density or the AS:SS ratio remaining unaltered. Indeed, an imbalance between inhibitory and excitatory activity has been previously described in the ventral tegmental area after repeated cocaine exposure^52,53^.

#### Postsynaptic targets

The most relevant change is an increase of 53% in the proportion of axodendritic AS on aspiny shafts —that in general originate from interneurons^54^— after cocaine-SA (it increased from 0.79% to 1.21% of the AS population). It has been previously observed that when long term potentiation (LTP) is induced exclusively in excitatory synapses on hippocampal pyramidal neurons, the temporal fidelity of synaptic integration and action potential generation in pyramidal cells is compromised. However, when LTP also occurs at excitatory synapses on feed-forward interneurons, temporal fidelity is preserved and information processing can be maintained without degradation during memory encoding^55^. It is possible that the increase in the proportion of AS on aspiny shafts after cocaine-SA has a relevant role in memory formation and consolidation induced by the drug.

#### Synaptic shape, synaptic size and curvature

Regarding the synaptic shape, we observed a higher proportion of synapses with more complex shapes — particularly in the case of axospinous AS and axodendritic SS (see **Supplementary table 8**). As stated above, synapses with more complex shapes are larger than macular ones; thus, the increase in the number of synapses with more complex shapes means an increase in synaptic surface.

We did not find significant differences in the mean SAS areas or perimeters, for either AS or SS after cocaine-SA (with the exception of an increase in the mean SAS perimeter of SS). However, there were differences in the frequency distribution of the SAS area and perimeter of synapses after cocaine-SA. Specifically, we observed a greater proportion of larger synapses, both AS and SS. For AS, this increase was mainly found within macular axospinous AS, while for SS it was not so clearly associated with any specific synaptic type. However, data regarding SS should be interpreted with caution since a lower number of synapses were analyzed compared to AS.

Taking both results together (regarding synaptic shape and size), cocaine-SA seems to induce an increase in the synaptic surface of axospinous AS, increasing both the proportion of synapses with more complex shapes and the size of macular synapses. In a previous study, we observed an increase in the proportion of larger dendritic spines in the Lewis rat CA1 after cocaine-SA^14^. We can therefore speculate that cocaine-SA causes an increase in synaptic activity and that this may induce the growth of macular synapses and, in some cases, lead to deep indentations and perforations appearing as the postsynaptic density (PSD) becomes larger (**Supplementary figure 2**).

Finally, several studies have analyzed the effect of cocaine-SA on the synaptic plasticity of the rat hippocampus in Lewis rats and other strains. LTP facilitation has been observed both after 3 weeks of cocaine-SA^18^ and following passive cocaine administration^17^. In addition, there is an impaired LTP depotentiation after 3 weeks of cocaine-SA, which seems to suggest a loss of synaptic flexibility^22^. Thus, it is possible that changes in the synaptic plasticity (LTP facilitation and impaired LTP depotentiation) and the results found in our study (larger synapses) after cocaine-SA may be related. In this regard, the synapse size has been associated with different functional attributes. For example, it has been proposed that small dendritic spines (with small synapses) are preferential sites for LTP induction, while large dendritic spines (with large synapses) may represent physical long-term memory storage locations^56,57,58^. Our data shows that synaptic size follows a unimodal and continuous distribution, so synapses cannot be divided into two groups based on their size. Therefore, if the function of the ‘learning or memory’ synapse depends on the synaptic size, there would be no clear transition between the two types of synapses. The strengthening of the synaptic transmission after cocaine-SA (inferred from the increase in the synaptic surface) could have various functional consequences with regard to electrically-induced synaptic plasticity (^56,57,^ see also^58^). One possibility is that this synaptic strengthening uses part of the capacity available in CA1 neurons to generate a greater synaptic plasticity, thus preventing the subsequent facilitation of electrically-induced LTP. Alternatively, it may be that stronger synapses more effectively facilitate LTP. Based on our results and the previous electrophysiological studies already mentioned, the second hypothesis seems to be more plausible. In addition, the increase in the size of SS after cocaine-SA could be the result of a compensatory mechanism to maintain the excitatory activity within certain limits^59^.

Our results suggest that the structural reorganization of synaptic circuits plays a role in their capacity for synaptic plasticity, which may be a regulatory factor of the actions that cocaine appears to have in the potentiation of context-dependent memories during cocaine-SA. Further studies will be necessary to evaluate the long-term effects of cocaine-SA during drug withdrawal periods, to have a more comprehensive picture of the synaptic alterations occurring in different phases of the transition from initial drug use to drug addiction.

## METHODS

### Animals

Adult male Lewis rats (n=13) were used in this study, weighing 275–350 g (Harlan Interfauna Ibérica, Barcelona, Spain) at the beginning of the experiments. They were housed individually in a climate-controlled room (23°C) with a 12h light–dark cycle (08:00–20:00 lights on) in accordance with the European Union guidelines for the care of laboratory animals (Directive 2010/63/EU) and we followed the “Principles of laboratory animal care”. All the animals were experimentally naïve and, unless otherwise specified, they had free access to Purina laboratory feed (Panlab, Barcelona, Spain) and tap water.

### Catheter Surgery and Cocaine Self-Administration procedure

The surgical procedure has been described elsewhere^13^. Briefly, rats were anaesthetized with a mixture of ketamine (40 mg/kg) and diazepam (10 mg/kg) and implanted with polyvinylchloride tubing (0.064 i.d.) into the jugular vein approximately at the level of the atrium. To evaluate the catheter patency and prevent infection, the catheters were flushed every day with 0.3 mL of an antibiotic solution (gentamicin, 0.10 mg/mL) dissolved in heparinized saline solution. Catheter patency was tested at the end of the experiments by infusing with thiopental (10 mg/kg, i.v.), and a good patency was assumed if the rat immediately lost consciousness.

Eighteen operant conditioning chambers (Coulburn Instruments, Allentown, PA, USA) with two fixed levers were used for the behavioral studies. To facilitate acquisition of cocaine self-administration, rats were food-deprived until they reached 95% of their free-feeding weight, and they were then placed on a fixed ratio (FR)-1 schedule of food reinforcement that was subsequently raised to FR-3 for several 30-min sessions. After lever-press training (rats showed a stable rate of lever pressing during the FR-3 schedule), the animals were fed ad libitum, and when animals recovered their free-feeding weight, catheter surgery was performed (see above). After a postoperative recovery period (7 days), the rats were food restricted —again to 95% of their original weight— and they were trained to respond for cocaine (n=7) or saline (n=6) infusions in daily 2-h sessions or until the rats earned 20 cocaine infusions for 21 days (7 d/week). A microliter injection pump (Harvard 22; Harvard Apparatus) was used to deliver intravenous cocaine (1 mg/kg) or saline infusions (100 µL) over 10 s, while inactive lever presses were recorded but had no programmed consequences. The FR ratio began on FR1 and was subsequently raised weekly, first to FR-2 and then to FR-3. A timeout period of 10 s was imposed in all the schedules. Following the 21 self-administration sessions, saline and cocaine-SA animals were intracardially perfused 24h after the last session (see details below).

Rats that self-administered cocaine progressively increased the number of infusions received and the number of times the lever was pressed with respect to the saline group (whose response was virtually null), reaching a stable self-administration pattern at the end of the 21 days. Animals that received cocaine showed more robust responses —number of lever presses— than saline animals during the last seven sessions under the FR-3 schedule of reinforcement (Saline: 2.14±1.19; Cocaine: 25.38±4.74; t_10_=11.65, p<0.0001). Total cocaine intake in SA animals was 161.85±26.24 mg/kg (average per session: 8.03±1.18 mg/kg) (Selvas et al., in preparation).

### Tissue preparation for electron microscopy

Rats were intracardially perfused with 100 ml of 1% paraformaldehyde in 0.1M phosphate buffer (PB; pH 7.3) followed by 500 ml of 4% paraformaldehyde and 0.125% glutaraldehyde in 0.1M PB (pH 7.3) at a flow rate of 25 ml/min. The descending aorta was clamped to preferentially restrict the flow of fixative to the upper body and accelerate perfusion of the brain. Immediately after perfusion, the brains were removed and postfixed for 6h at room temperature in the same fixative. After postfixation, 150 µm coronal vibratome (Lancer 1000; St Louis, MO, USA) sections were obtained.

Selected sections containing the dorsal hippocampus were washed in sodium cacodylate (Sigma, C0250-500G, Germany) buffer (0.1M) and postfixed for 24h in a solution containing 2% paraformaldehyde, 2.5% glutaraldehyde (TAAB, G002, UK) and 0.003% CaCl_2_ (Sigma, C-2661-500G, Germany) in sodium cacodylate buffer (0.1M). These sections were washed in sodium cacodylate buffer (0.1M) and treated with 1% OsO_4_ (Sigma, O5500, Germany), 0.1% ferrocyanide potassium (Probus, 23345, Spain), 0.003% CaCl_2_ and 7% glucose in sodium cacodylate buffer (0.1M) for 1h at room temperature. After washing in sodium cacodylate buffer (0.1M), the sections were stained with 2% uranyl acetate (EMS, 8473, USA), and then dehydrated and flat-embedded in Araldite (TAAB, E021, UK) for 48h at 60°C^60^. Embedded sections were glued onto a blank Araldite block and trimmed. Semithin sections (1–2 µm thickness) were obtained from the surface of the block until the region of interest was reached. Then, semithin sections were stained with 1% toluidine blue (Merck, 115930, Germany) in 1% sodium borate (Panreac, 141644, Spain). The last semithin section (which corresponds to the section immediately adjacent to the block surface) was examined under light microscope and photographed to accurately locate the region to be examined.

The blocks containing the embedded tissue were glued onto a sample stub using conductive adhesive tabs (EMS 77825-09, Hatfield, PA, USA). All the surfaces of the block —except for the one to be studied (the top surface)— were covered with silver paint (EMS 12630, Hatfield, PA, USA) to prevent charging artifacts. The stubs with the mounted blocks were then placed into a sputter coater (Emitech K575X, Quorum Emitech, Ashford, Kent, UK) and the top surface was coated with a 10–20 nm thick layer of gold/palladium to facilitate charge dissipation.

### Focused ion beam milling and the acquisition of serial scanning electron microscopy images

We used a Crossbeam 540 electron microscope (Carl Zeiss NTS GmbH, Oberkochen, Germany) as described in ^61^. This instrument combines a high-resolution field emission SEM column with a focused gallium ion beam, which can mill the sample surface, removing thin layers of material on a nanometer scale. After removing each slice (20 nm thick), the milling process was paused, and the freshly exposed surface was imaged with a 1.8-kV acceleration potential using the in-column energy selective backscattered (EsB) electron detector. The milling and imaging processes were sequentially repeated, and long series of images were acquired through a fully automated procedure, thus obtaining a stack of images that represented a three-dimensional sample of the tissue^61^.

Twenty-four different samples (stacks of images) were acquired in the neuropil of the *stratum radiatum* —at a distance of between 70 and 100 µm from the border of the *stratum pyramidale*— from the CA1 hippocampal region in coronal slices of 12 Lewis rats (6 from the saline group and 6 from the cocaine-SA group; 2 stacks of images were taken per animal) (**Figure 1**). Image resolution in the x,y plane was 5 nm / pixel. Resolution in the z-axis (section thickness) was 20 nm and image sizes were 2048 × 1536 pixels. The number of sections per stack ranged from 210 to 360 (mean 265.29; total 6,412 sections) (**Figure 2 and Supplementary figure 1**).

### Tissue shrinkage estimation

All measurements were corrected for tissue shrinkage that occurs during osmication and plastic embedding of the vibratome sections containing the area of interest as described by ^61^. We measured the surface area and thickness of the vibratome sections with Stereo Investigator (MBF Bioscience, Williston, VT, USA), both before and after they were processed for electron microscopy^62^. The surface area after processing was divided by the value before processing to obtain an area shrinkage factor (p^2^) of 0.97. The linear shrinkage factor for measurements in the plane of section (p) was therefore 0.99. The shrinkage factor in the z-axis was 0.96. All distances measured were corrected to obtain an estimate of the pre-processing values.

### 3D analysis of synapses

#### Classification of synapses, identification of the postsynaptic target, the synaptic shape and morphological measurements

Stacks of images obtained by the FIB/SEM were analyzed using EspINA software (EspINA Interactive Neuron Analyzer, 2.8.2; https://cajalbbp.es/espina/), which allows the segmentation of synapses in stacks of serial sections (for a detailed description of the segmentation algorithm, see^63^; **Figure 3**). Since the synaptic junctions were fully reconstructed as described elsewhere^61^, each synapse could be classified as AS or SS based on its prominent or thin postsynaptic density (PSD), respectively (**Figure 2 and Supplementary figure 1**; ^64,65^). EspINA provides the number of synapses within an unbiased 3D counting frame (CF) of known volume, so the local density of synapses could be established (**Figure 3c**; ^66^ (see^61^ for details)).

The 3D segmentation of synaptic junctions includes both the presynaptic density (active zone; AZ) and the PSD. Since the AZ and the PSD are located face to face, their surface areas are very similar (correlation coefficients over 0.97; ^67,68^). Thus, they can be simplified to a single surface and represented as the surface of apposition between the AZ and the PSD. This surface can be extracted from the 3D segmented synaptic junction^69^. For the sake of clarity, we have referred to this surface as the synaptic apposition surface (SAS) (**Figure 3d–i**). The SAS areas and perimeters of each synaptic junction were extracted with EspINA software to study morphological parameters regarding synapses. This software also permits the quantitation of the curvature of the synapses as it adapts to the curvature of the synaptic junction. Specifically, curvature measurements are calculated as 1 minus the ratio between the projected area of the SAS and the area of the SAS^63^. This measurement would be 0 in a flat SAS and would increase to a maximum value of 1 as the SAS curvature increases.

Additionally, based on the postsynaptic targets, synapses were further classified as axospinous synapses (synapses on dendritic spines) and axodendritic synapses (synapses on dendritic shafts). In the case of axospinous synapses, they were further subdivided into axospinous synapses on the head or on the neck of the spine. For axodendritic synapses, dendritic shafts were further classified as spiny (when dendritic spines could be observed emerging from the shaft) or aspiny. As described in ^24^, only clearly identifiable postsynaptic elements were quantified (i.e., elements that were unambiguously identified from navigating through the stack of images; **Figure 2**).

Finally, synapses were classified —according to the shape of their synaptic junction— into four categories, as described elsewhere^46^. Synapses with a flat, disk-shaped PSD were classified as macular. A second category was established by the presence of an indentation in the perimeter (horseshoe-shaped synapses). Synaptic junctions with one or more holes in the PSD were referred to as perforated. Synaptic junctions with two or more physically discontinuous PSDs were categorized as fragmented (**Figure 3f–i**, see **Figure 6a**).

#### Spatial distribution analysis of synapses

To analyze the spatial distribution of synapses, Spatial Point Pattern analysis was performed as described elsewhere^70,41^. Briefly, we recorded the spatial coordinates of the centers of gravity or centroids of synaptic junctions in 24 samples obtained from the corresponding stacks of serial sections. We then calculated the F, G and K functions to compare the actual position of centroids of synapses with the Complete Spatial Randomness (CSR) model, or homogeneous spatial Poisson point process, where a point is equally likely to occur at any location within a given volume. The F function or empty-space function is the cumulative plot of distances between a regular grid of points and the closest sample points. The G function is the cumulative plot of distances from each point to its nearest neighbor. The K function is given by the number of points within a sphere of increasing radius, centered on each sample point^71,72^. We used R software (https://www.r-project.org/) and spatstat package^73,74^ (http://spatstat.org/) for the calculations. A sample was considered to be compatible with the CSR model when the observed F, G and K functions lay within the envelope generated by 99 simulations of the CSR model (see **Figure 4c**). Deviations in G functions occur because synaptic junctions cannot overlap, and thus the minimum intersynaptic distances (measured between their centroids) must be limited by the sizes of the synaptic junctions themselves (see **Figure 4c** and ^41^).

### Estimation of the volume fraction occupied by mitochondria

See Supplementary Methods for information about the estimation of the volume fraction occupied by mitochondria.

### Statistical analysis

Statistical analysis of the data was carried out using GraphPad Prism statistical package (Prism 7.00 for Windows, GraphPad Software Inc., USA), SPSS software (IBM SPSS Statistics for Windows, Version 24.0. Armonk, NY: IBM Corp) and R Project software (R 3.5.1; Bell Laboratories, NJ, USA; http://www.R-project.org). Unpaired parametric tests were used when normality and homoscedasticity criteria were met; otherwise, equivalent non-parametric tests were used. Differences in (i) the mean synaptic density, (ii) the mean SAS area, SAS perimeter or SAS curvature, and (iii) the mean volume fraction occupied by mitochondria were analyzed by performing a two-sided, t-Student test or one-way analysis of variance (ANOVA) (with Bonferroni correction post-hoc tests). When necessary, the equivalent non-parametric Mann-Whitney U (MW) test or Kruskal-Wallis (KW) test (with MW pairwise comparisons) was used, respectively. In addition, when comparing mean values, the “n” per group corresponds to the number of animals. Frequency distributions were analyzed using Kolmogorov-Smirnov (KS) nonparametric tests. To perform statistical comparisons of AS and SS, postsynaptic targets and synaptic shapes proportions, χ^2^ test was used for contingency tables. In some χ^2^ statistical analyses, we firstly performed an “omnibus test” based on 2 × 4 contingency tables. To further investigate the specific cells driving the significance of the χ^2^ test, a partitioning procedure was applied to create 2 × 2 contingency tables^75^. Note that due to the large sample size of synapses analyzed —especially for AS whose numbers were in the thousands— statistical differences in KS and χ^2^ tests should be carefully considered when p>0.001 to avoid the overestimation of differences.

## ACKNOWLEDGMENTES

We would like to thank Rosa Ferrado, Carmen Álvarez, Miriam Martín and Lorena Valdés for their helpful technical assistance and Nick Guthrie for his excellent text editing.

This work was supported by grants from the following entities: Ministerio de Ciencia e Innovación (Reference number: PSI2016-80541-P); Ministerio de Sanidad, Servicios Sociales e Igualdad (Red de Trastornos Adictivos — Reference number RTA-RD16/0017/0022 del Instituto de Salud Carlos III— and Plan Nacional sobre Drogas, Reference number: 2016I073); UNED (Plan de Promoción de la Investigación, 2014-040-UNED-POST); UNED-Banco Santander (“Independent Thinking”, reference number: 2017-VICE-0012); Unión Europea (Reference number: JUST-2017-AG-DRUGS-806996-JUSTSO); Centro de Investigación en Red sobre Enfermedades Neurodegenerativas (CIBERNED, CB06/05/0066, Spain); the Spanish “Ministerio de Ciencia, Innovación y Universidades” (grant PGC2018-094307-B-I00 and the Cajal Blue Brain Project [the Spanish partner of the Blue Brain Project initiative from EPFL, Switzerland]); and the European Union’s Horizon 2020 Research and Innovation Programme under grant agreement No. 785907 (Human Brain Project, SGA2). MM-C was awarded a research fellowship from the Spanish Ministry of Education, Culture and Sports (contract FPU14/02245).

## AUTHOR CONTRIBUTIONS

EA, JDeF, LB-L and MM oversaw and designed the project. MM and LB-L designed and performed experiments and data analysis. AS, JG-S and MM-C performed experiments and helped to interpret experiments. AS and MM performed catheter surgery and the cocaine-SA procedure. JDeF and L-BL wrote the manuscript. EA and JDeF provided financial and intellectual support to the research. All authors read, reviewed, helped to edit, and approved the final manuscript.

## DATA AVAILABILITY

The datasets generated and/or analyzed during the current study are available from the corresponding author on reasonable request.

