## Supplementary Material for "3D SYNAPTIC ORGANIZATION OF THE RAT CA1 AND ALTERATIONS INDUCED BY COCAINE SELF-ADMINISTRATION"

### SUPPLEMENTARY TEXT: METHODS, RESULTS AND DISCUSSION

#### Distribution of postsynaptic targets

*Results*

We also determined the proportions of single and multiple synapses per spine head. The majority of synapses were single AS (99.28%). Different combinations of multiple synapses on a spine head were found as follows: 0.11% had two AS and 0.61% had one AS and one SS (**Supplementary table 9**).

No differences in the number of multiple synapses were found after cocaine-SA ($\chi^2$=0.254, p=0.614) (**Supplementary table 9**).

*Discussion*

The percentage of dendritic spines establishing multiple synapses was lower (0.72%) than reported in the somatosensory cortex of the rat (5.57%), the human CA1 (2.11%) or in layer II of the human transentorhinal cortex (5.52%)[1,2,3].

#### The synaptic apposition surface (SAS)

***SAS area and perimeter (synaptic size)***

*Results*

Additionally, we studied the synaptic size with regard to the postsynaptic targets. For AS, the mean SAS area and perimeter of axospinous AS were not significantly different to those of axodendritic AS ($t_{10}$=-0.558, p=0.596 for area and $t_{10}$=-1.016, p=0.336 for perimeter; **Figure 7c**; **Supplementary table 6**). For SS, axospinous SS were significantly smaller than axodendritic SS ($t_{10}$=-4.256, p=0.002 for area and $t_{10}$=-3.635, p=0.005 for perimeter; **Figure 7c**; **Supplementary table 6**). Regarding the type of dendritic shaft, we observed that synapses on aspiny shafts were larger than synapses on spiny shafts, both for AS ($t_8$=-2.383, p=0.044 for area and $t_8$=-2.342, p=0.047 for perimeter) and SS, but in this case the difference was only statistically significant for the SAS area ($t_8$=-2.537, p=0.039 for area and $t_{10}$=-1.458, p=0.270 for perimeter) (**Figure 7d; Supplementary table 6**).

Finally, we analyzed differences in the synaptic size according to the shape of the synaptic junctions (**Figure 7e; Supplementary table 7**). Within AS, statistically significant differences were found in the mean SAS area and perimeter of synapses classified according to their shape (KW=14.790, p=0.002 for area; KW=19.238, p=0.000 for perimeter). Specifically, macular AS had a smaller area and perimeter than fragmented AS (MW=-14.5, p=0.002 for area and MW=-15.9, p=0.001 for perimeter), a smaller area than perforated AS (MW=-11.833, p=0.015) and a smaller perimeter than horseshoe-shaped AS (MW=-13.333, p=0.004) (**Figure 7e**; **Supplementary table 7)**. Within SS, statistically significant differences were also found in the mean SAS area and perimeter of synapses classified according to their shape (KW=11.591, p=0.003 for area and KW=12.877, p=0.002 for perimeter). Specifically, macular SS were smaller than horseshoe-shaped SS (MW=-10.167, p=0.003 for area and MW=-11, p=0.001 for perimeter; **Figure 7e**; **Supplementary table 7**).

*Discussion*

Similar to observations in the human CA1, axodendritic AS in the rat CA1 were a very similar size to axospinous AS, but we observed axodendritic SS to be larger than axospinous SS[3]. Furthermore, similar to the case of the rat somatosensory cortex and the human CA1[4,3], we found a correlation between the synaptic area and perimeter for AS and SS. Therefore, the larger the SAS area of a synapse, the more tortuous its perimeter — the SAS perimeter tends to grow faster than the perimeter of a circle.

***SAS curvature***

*Results*

*Saline group*

Curvature measurements of all synapses, AS and SS had a slightly inverted association to area ($R^2$=-0.332, p=0.000 for all synapses; $R^2$=-0.325, p=0.000 for AS; $R^2$=-0.457, p=0.000 for SS) and perimeter ($R^2$=-0.301, p=0.000 for all synapses; $R^2$=-0.295, p=0.000 for AS; $R^2$=-0.401, p=0.000 for SS).

Symmetric synapses were observed to be flatter than AS (mean SAS curvature of AS: 0.0037; mean SAS curvature of SS: 0.0028) (MW=5, p=0.041) (**Table 1**).

The frequency histograms of SAS curvature ratios showed a positive skewness with larger number of synapses presenting lower values, meaning a larger prevalence of flatter synapses than more curved ones for both AS and SS populations. The SAS curvature distribution of AS statistically differed from SS (KS=2.092, p=0.000) and a longer right tail was observed for AS.

With regard to the postsynaptic targets, the mean curvature ratio of axospinous and axodendritic synapses did not differ for either AS or SS (MW=12, p=0.662 for AS and MW=21, p=0.699 for SS) (**Supplementary table 6**).

Regarding the synaptic shape, we did not find differences in the mean curvature ratio within AS (KW=6.677, p=0.083), while differences were found within SS ($F_{2,15}$=10.466, p=0.001) — macular SS were flatter than horseshoe-shaped SS and perforated SS ($t_{15}$=4.317, p=0.002 and $t_{15}$=3.439, p=0.011, respectively) (**Supplementary table 7**).

*Cocaine-SA group*

No differences were found in the mean curvature ratio of AS and SS after cocaine-SA ($t_{10}$=0.838, p=0.422 for AS, MW=16, p=0.818 for SS) (**Table 1**). Likewise, after cocaine-SA, we did not find significant differences in the mean SAS area or perimeter of AS and SS classified according to the postsynaptic targets or the synaptic shapes (**Supplementary tables 6, 7**).

*Discussion*

SS were observed to be flatter than AS, which was different from observations made in our previous studies in the human CA1 and transentorhinal cortex (in which no differences were found) and also different from observations in the rat somatosensory cortex (where AS were flatter than SS)[5,4,3]. As previously discussed in Santuy et al.[4], changes in synaptic curvature have been associated with synaptic efficacy. However, unlike synaptic area, it is still not known how curvature influences synaptic functioning. Furthermore, unlike Santuy et al.[4] and Montero-Crespo et al.[3], who did not find a correlation between synaptic size and curvature, we found a slight inverted correlation for synaptic size and curvature in the present study. Thus, more research is necessary to elucidate the biological relevance of this morphological parameter.

**Volume fraction of neuropil occupied by mitochondria**

*Methods*

We used the Cavalieri method[6] for the estimation of the volume fraction of neuropil occupied by mitochondria in the stacks of FIB/SEM images. We used "Image J Stereology Toolset"[7] to analyze the 24 stacks of images described above. A grid with an area per point of 0.25 µm$^2$ was used. The estimations were made in every 10th section of each stack (z=200nm). A total of 624 sections were analyzed. The parameters used for the Cavalieri method (grid size and number of sections) were chosen based on a pilot study[8]. Coefficients of error and variation were calculated to ensure the reliability of the measurements[9].

*Results*

Mitochondria play a key role in energy production and calcium buffering, among many other functions, and a positive correlation between the volume fraction of mitochondria located in neuronal processes and the density of synapses has been found[10]. Thus, we evaluated possible changes in the volume fraction of neuropil occupied by mitochondria ($V_m$) in the *stratum radiatum* of the CA1 field after cocaine-SA. No differences were found ($V_m$=7.31 % in saline group and $V_m$=7.49 % in cocaine-SA group; $t_{10}$=-0.601, p=0.561) (**Table 1**).

*Discussion*

We did not observe that the synaptic size increase was accompanied by changes in the volume fraction of the neuropil occupied by mitochondria. This is interesting because mitochondria are described to be involved in the morphogenesis and plasticity of dendritic spines and synapses[11,12]. Thus, further studies about the possible relationship between mitochondria, synaptic organization and cocaine effects should be performed.

**SUPPLEMENTARY FIGURES**

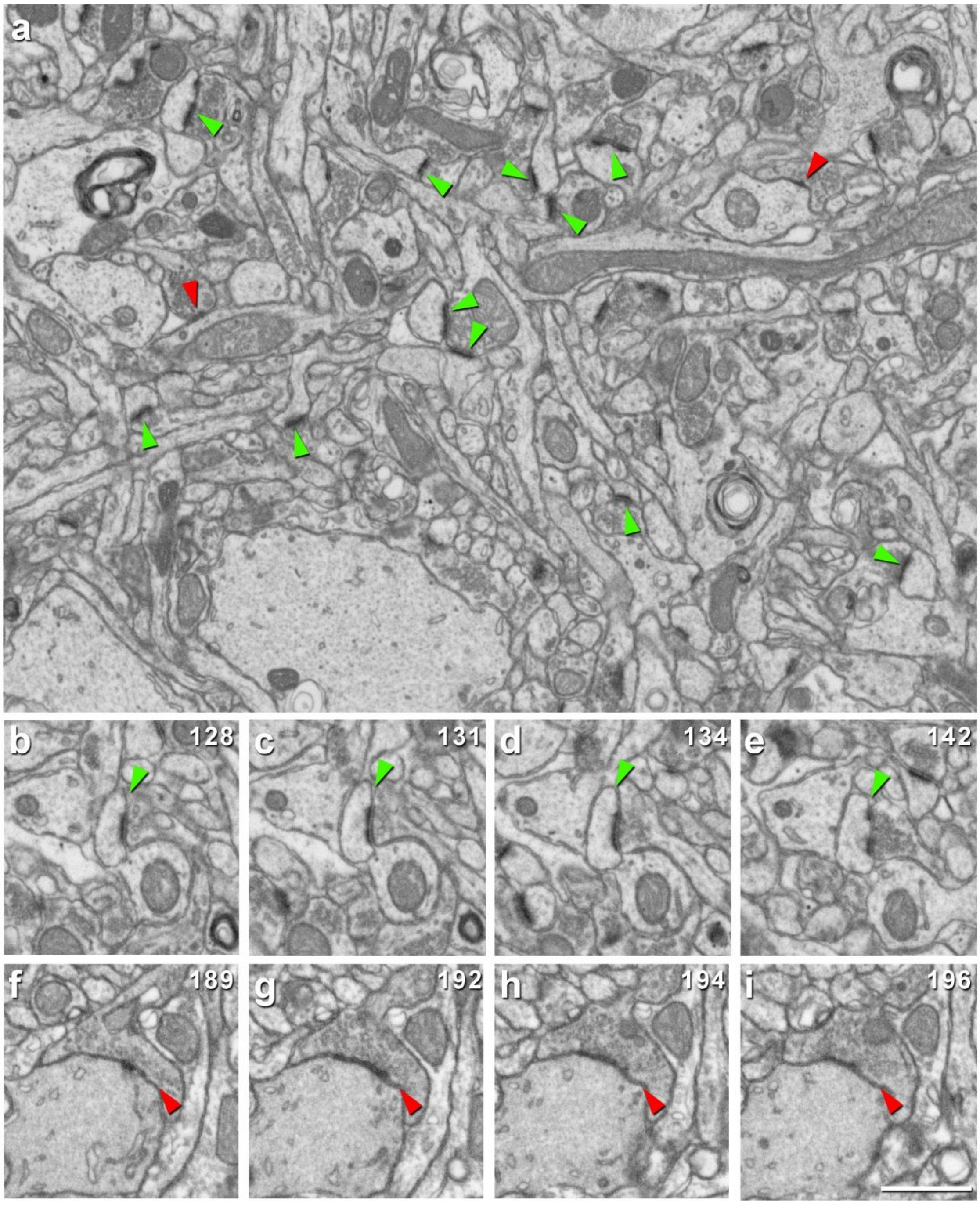


**Supplementary figure 1. FIB/SEM images in the *stratum radiatum* of the rat CA1**. **a**, Example of an FIB/SEM image from a stack of images. Some asymmetric and symmetric synapses have been pointed out (green and red arrowheads, respectively). **b**–**e**, Same asymmetric synapse in different sections from the stack of images. **f**–**i**, Same symmetric synapse in different sections from the stack of images. The number of the section is indicated in the top right-hand corner of each section. Scale bar in **i** corresponds to: 0.95 µm in **a**, 0.79 µm in **b**–**i**.

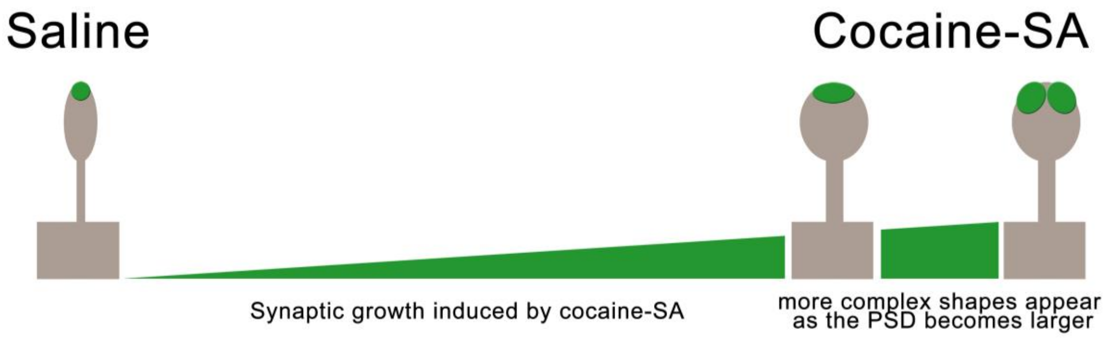


**Supplementary figure 2**. **Summary of the main synaptic alterations induced by cocaine-SA.** We can speculate that cocaine-SA causes an increased synaptic activity that induces the growth of macular asymmetric synapses and, in some cases, deep indentations and perforations appear as the PSD becomes larger. See **Supplementary table 8** for more detailed information. PSD: postsynaptic density; SA: self-administration.

SUPPLEMENTARY TABLES

| Group | Type of synapse | Dendritic spine | Spine heads | Spine necks | Dendritic shaft | Spiny dendritic shaft | Aspiny dendritic shaft | Total synapses |
|---|---|---|---|---|---|---|---|---|
| Saline | AS | 98.34% (4738) | 98.32% (4737) | 0.02% (1) | 1.66% (80) | 0.87% (42) | 0.79% (38) | 100% (4818) |
| | SS | 17.53% (37) | 14.69% (31) | 2.84% (6) | 82.47% (174) | 77.73% (164) | 4.74% (10) | 100% (211) |
| Cocaine -SA | AS | 97.73% (4432) | 97.62%* (4427) | 0.11% (5) | 2.27% (103) | 1.06% (48) | 1.21%* (55) | 100% (4535) |
| | SS | 16.76% (29) | 14.45% (25) | 2.31% (4) | 83.24% (144) | 78.61% (136) | 4.63% (8) | 100% (173) |

**Supplementary table 1. Distribution of asymmetric (AS) and symmetric (SS) synapses on spines and dendritic shafts in the *stratum radiatum* of the rat CA1 in the saline and the cocaine-SA groups**. Synapses on spines have been sub-divided into those that are established on spine heads and those that are established on spine necks. Moreover, we differentiated between spiny and aspiny dendritic shafts. Data are expressed as percentages, and absolute numbers of synapses studied are given in parentheses. Asterisks point out those synapses whose proportion varied after cocaine-SA: a reduction in the proportion of AS on spine heads and an increase in the proportion of AS on aspiny shafts (see text for further information). AS: asymmetric synapses; SS: symmetric synapses.

| | | Postsynaptic target | | | | |
|---|---|---|---|---|---|---|
| | | Spine Head | Spine Neck | Spiny Shaft | Aspiny Shaft | Total |
| Type of Synapse | AS | **4737** (4568) | **1** (6.7) | **42** (197.4) | **38** (46) | 4818 |
| | SS | **31** (200) | **6** (0.3) | **164** (8.6) | **10** (2) | 211 |
| | Total | 4768 | 7 | 206 | 48 | 5029 |

**Supplementary table 2**. An example of a 2 × 4 contingency table showing the type of synapse (AS: asymmetric synapse; SS: symmetric synapse) against the postsynaptic target (Spine Head, Spine Neck, Spiny Shaft, Aspiny Shaft). The bold numbers indicate an observed value, while the numbers in brackets are expected values. $\chi^2$=3212.684, p=0.000. To determine which values in this test are driving this significance, 2 × 2 contingency tables were created (see **Supplementary table 3**)

| | | Postsynaptic target | | |
|---|---|---|---|---|
| | | Spine Head | Non-Spine Head | Total |
| **Type of Synapse** | AS | **4737** (4568) | **81** (250) | 4818 |
| | SS | **31** (200) | **180** (11) | 211 |
| | Total | 4768 | 261 | 5029 |

| | | Postsynaptic target | | |
|---|---|---|---|---|
| | | Spine Neck | Non-Spine Neck | Total |
| **Type of Synapse** | AS | **1** (6.7) | **4817** (4811.3) | 4818 |
| | SS | **6** (0.3) | **205** (210.7) | 211 |
| | Total | 7 | 5022 | 5029 |

| | | Postsynaptic target | | |
|---|---|---|---|---|
| | | Spiny Shaft | Non-Spiny Shaft | Total |
| **Type of Synapse** | AS | **42** (197.4) | **4776** (4620.6) | 4818 |
| | SS | **164** (8.6) | **47** (202.4) | 211 |
| | Total | 206 | 4823 | 5029 |

| | | Postsynaptic target | | |
|---|---|---|---|---|
| | | Aspiny Shaft | Non-Aspiny Shaft | Total |
| **Type of Synapse** | AS | **38** (46) | **4780** (4772) | 4818 |
| | SS | **10** (2) | **201** (209) | 211 |
| | Total | 48 | 4981 | 5029 |

**Supplementary table 3**. Example of the 2 × 2 contingency tables to evaluate which values in the 2 × 4 contingency table in **Supplementary table 2** are driving the significant difference. $\chi^2$=2873.067, p=0.000; $\chi^2$=115.886, p=0.000; $\chi^2$=3039.292, p=0.000; $\chi^2$=33.374, p=0.000, respectively.

| Group | Type of synapse | Macular synapses | Horseshoe -shaped synapses | Perforated synapses | Fragmented synapses | Total synapses |
|---|---|---|---|---|---|---|
| **Saline** | AS | 88.23% (4325) | 4.24% (208) | 7.16% (351) | 0.37% (18) | 100% (4902) |
| | SS | 86.51% (186) | 4.65% (10) | 8.84% (19) | 0.0% (0) | 100% (215) |
| **Cocaine-SA** | AS | 86.60% (3974) | 4.49% (206) | 8.02% (368) | 0.89% (41) | 100% (4589) |
| | SS | 74.16% (132) | 12.36% (22) | 12.92% (23) | 0.56% (1) | 100% (178) |

**Supplementary table 4. Proportion of the different synaptic shapes in the *stratum radiatum* of the rat CA1 in the saline and the cocaine-SA group**. Data are given as percentages; absolute number of synapses studied is indicated in parentheses. AS: Asymmetric synapses; SS: symmetric synapses.

Statistical comparisons of the saline group *versus* the cocaine-SA group (see **Supplementary tables 2 and 3** for an example of the analysis performed):

- Contingency tables 2 x 4 (saline-cocaine x synaptic shapes):

For AS: $\chi^2$=13.916, p=0.003
For SS: $\chi^2$=11.531, p=0.005

- Contingency tables 2 x 2:

For AS:
saline-cocaine x macular-non macular: $\chi^2$=5.741, p=0.017
saline-cocaine x HS-non HS: $\chi^2$=0.343, p=0.558
saline-cocaine x perforated-non perforated: $\chi^2$=2.497, p=0.114
saline-cocaine x fragmented-non fragmented: $\chi^2$=10.625, p=0.001
For SS:
saline-cocaine x macular-non macular: $\chi^2$=9.625, p=0.002
saline-cocaine x HS-non HS: $\chi^2$=7.736, p=0.005
saline-cocaine x perforated-non perforated: $\chi^2$=1.702, p=0.192

| Group | Type of synapse | Postsynaptic target | Macular synapses | Horseshoe-shaped synapses | Perforated synapses | Fragmented synapses | Total synapses |
|---|---|---|---|---|---|---|---|
| **Saline** | AS | Axospinous | 87.99% (4169) | 4.33% (205) | 7.30% (346) | 0.38% (18) | 100% (4738) |
| | | Axodendritic | 95% (76) | 2.5% (2) | 2.5% (2) | 0% (0) | 100% (80) |
| | SS | Axospinous | 94.59% (35) | 0% (0) | 5.41% (2) | 0% (0) | 100% (37) |
| | | Axodendritic | 84.48% (147) | 5.75% (10) | 9.77% (17) | 0.0% (0) | 100% (174) |
| **Cocaine-SA** | AS | Axospinous | 86.15% (3819) | 4.62% (205) | 8.30% (368) | 0.93% (40) | 100% (4432) |
| | | Axodendritic | 99.02% (101) | 0.98% (1) | 0% (0) | 0% (0) | 100% (102) |
| | SS | Axospinous | 93.10% (27) | 6.90% (2) | 0% (0) | 0% (0) | 100% (29) |
| | | Axodendritic | 71.53% (103) | 13.19% (19) | 14.58% (21) | 0.70% (1) | 100% (144) |

**Supplementary table 5. Proportion of the different synaptic shapes depending on the type of synapse and the postsynaptic target in the *stratum radiatum* of CA1 in the saline and cocaine-SA groups**. Data are given as percentages; absolute numbers of synapses studied are indicated in parentheses. AS: Asymmetric synapses; SS: symmetric synapses.

Statistical comparisons of the saline *versus* the cocaine-SA group (see **Supplementary tables 2 and 3** for an example of the analysis performed):

- Contingency tables 2 x 4 (saline-cocaine x synaptic shapes):

For axospinous AS: $\chi^2$=14.812, p=0.002
For axodendritic AS: $\chi^2$=2.940, p=0.232
For axospinous SS: $\chi^2$=3.347, p=0.158
For axodendritic SS: $\chi^2$=9.210, p=0.027

- Contingency tables 2 x 2:

For axospinous AS:
saline-cocaine x macular-non macular: $\chi^2$=6.871, p=0.009
saline-cocaine x HS-non HS: $\chi^2$=0.469, p=0.493
saline-cocaine x perforated-non perforated: $\chi^2$=3.161, p=0.075
saline-cocaine x fragmented-non fragmented: $\chi^2$=10.631, p=0.001
For axodendritic SS:

saline-cocaine x macular-non macular: $\chi^2=7.866$, $p=0.004$
saline-cocaine x HS-non HS: $\chi^2=5.273$, $p=0.018$
saline-cocaine x perforated-non perforated: $\chi^2=1.735$, $p=0.188$

| Group | Postsynaptic target | Type of synapse | Area of SAS (nm$^2$; mean± SEM) | Perimeter of SAS (nm; mean± SEM) | Curvature of SAS (mean± SEM) |
|---|---|---|---|---|---|
| Saline | Axospinous | AS | 42255±1960 | 989±28 | 0.0041±0.0005 |
| | | SS | 46001±5149 | 1137±94 | 0.0038±0.0016 |
| | Spine Heads | AS | 42263±1955 | 1009±25 | 0.0039±0.0005 |
| | | SS | 48775±6210 | 1185±115 | 0.0041±0.0020 |
| | Spine Necks | AS | 8004±0 | 363±0 | 0.0021±0 |
| | | SS | 25695±9050 | 761±165 | 0.0022±0.0004 |
| | Axodendritic | AS | 45403±5290 | 1075±87 | 0.0046±0.0015 |
| | | SS | 72331±3429 | 1554±65 | 0.0027±0.0002 |
| | Spiny Dendrite | AS | 31307±8212 | 805±136 | 0.0036±0.0004 |
| | | SS | 68000±5328 | 1496±94 | 0.0022±0.0005 |
| | Aspiny Dendrite | AS | 59613±7554 | 1230±80 | 0.0051±0.0030 |
| | | SS | 119876±28875 | 2090±396 | 0.0025±0.0010 |
| Cocaine-SA | Axospinous | AS | 48785±4534 | 1065±43 | 0.0038±0.0003 |
| | | SS | 41916±6555 | 1133±123 | 0.0029±0.0007 |
| | Spine Heads | AS | 48808±4542 | 1065±43 | 0.0036±0.0004 |
| | | SS | 42293±7279 | 1136±125 | 0.0032±0.0010 |
| | Spine Necks | AS | 29725±8882 | 861±183 | 0.0023±0.0011 |
| | | SS | 35078±1894 | 1110±224 | 0.0024±0.0011 |
| | Axodendritic | AS | 55858±13524 | 1143±180 | 0.0025±0.0003 |
| | | SS | 96559±16379 | 1958±240 | 0.0042±0.0019 |
| | Spiny Dendrite | AS | 54072±17828 | 1158±277 | 0.0025±0.0004 |
| | | SS | 97417±17342 | 1971±257 | 0.0042±0.0019 |
| | Aspiny Dendrite | AS | 75585±12674 | 1282±142 | 0.0021±0.0004 |
| | | SS | 89380±9819 | 1856±168 | 0.0018±0.0002 |

**Supplementary table 6. Data regarding size (area and perimeter) and curvature of the SAS from synapses on spines and dendritic shafts in *stratum radiatum* of the**

**rat CA1 in the saline and cocaine-SA groups.** AS: asymmetric synapses; SAS: synaptic apposition surface; SEM: standard error of the mean; SS: symmetric synapses.

Statistical comparisons of the saline group *versus* the cocaine-SA group:

AS axospinous: $t_{10}$=-1.322, p=0.216 for area, $t_{10}$=-1.467, p=0.173 for perimeter and $t_{10}$=0.019, p=0.985 for curvature; AS axodendritic: $t_{10}$=-0.720, p=0.497 for area, $t_9$=-0.316, p=0.759 for perimeter and MW=8, p=0.247 for curvature; SS axospinous: $t_{10}$=-0.490, p=0.635 for area, $t_{10}$=-0.024, p=0.981 for perimeter and MW=21, p=0.699 for curvature; SS axodendritic: $t_{10}$=-1.448, p=0.203 for area, $t_{10}$=-1.626, p=0.135 for perimeter and MW=17, p=0.937 for curvature.

| Group | Synaptic shape | Type of synapse | Area of SAS ($nm^2$; mean± SEM) | Perimeter of SAS (nm; mean± SEM) | Curvature of SAS (mean± SEM) |
|---|---|---|---|---|---|
| **Saline** | **Macular** | AS | 34962±1734 | 847±21 | 0.0037±0.0003 |
| | | SS | 56566±3770 | 1302±55 | 0.0030±0.0003 |
| | **HS** | AS | 89922±5155 | 2274±117 | 0.0020±0.0001 |
| | | SS | 183060±44439 | 3628±592 | 0.0013±0.0003 |
| | **Perforated** | AS | 97734±4756 | 1762±57 | 0.0089±0.0065 |
| | | SS | 117542±20127 | 2147±288 | 0.0016±0.0003 |
| | **Fragmented** | AS | 105173±8823 | 2545±152 | 0.0022±0.0005 |
| | | SS | - | - | - |
| **Cocaine -SA** | **Macular** | AS | 39713±4132 | 900±46 | 0.0039±0.0004 |
| | | SS | 62857±8891 | 1420±113 | 0.0052±0.0028 |
| | **HS** | AS | 111949±11303 | 2530±121 | 0.0018±0.0003 |
| | | SS | 131902±27987 | 2879±327 | 0.0025±0.0008 |
| | **Perforated** | AS | 105655±10147 | 1843±80 | 0.0017±0.0001 |
| | | SS | 186253±35546 | 3031±541 | 0.0013±0.0001 |
| | **Fragmented** | AS | 108308±11840 | 2681±239 | 0.0020±0.0003 |
| | | SS | 183837±0 | 5710±0 | 0.0008±0 |

**Supplementary table 7. Data regarding size (area and perimeter) and curvature of the SAS from synapses with different shapes in the *stratum radiatum* of the rat CA1 in the saline and cocaine-SA groups.** AS: asymmetric synapses; SAS: synaptic apposition surface; SEM: standard error of the mean; SS: symmetric synapses.

Statistical comparisons of the saline group *versus* the cocaine-SA group:

AS macular: $t_{10}$=-1.060, p=0.326 for area, $t_{10}$=-1.047, p=0.320 for perimeter and $t_{10}$=-0.285, p=0.781 for curvature; AS HS: $t_{10}$=-1.773, p=0.107 for area, $t_{10}$=-1.520, p=0.159 for perimeter and $t_{10}$=-0.944, p=0.368 for curvature; AS perforated: $t_{10}$=-0.707, p=0.496 for area, $t_{10}$=-0.826, p=0.428 for perimeter and MW=10, p=0.240 for curvature; AS fragmented: $t_9$=-0.205, p=0.842 for area, $t_9$=-0.455, p=0.660 for perimeter and $t_{10}$=-0.337, p=0.744 for curvature; SS macular: $t_{10}$=-0.651, p=0.529 for area, $t_{10}$=-0.944, p=0.376 for perimeter and MW=16, p=0.749 for curvature; SS HS: MW=11, p=0.310 for area, $t_{10}$=-1.520, p=0.159 for perimeter and MW=26, p=0.240 for curvature; SS perforated: $t_{10}$=-1.682, p=0.123 for area, $t_{10}$=-1.108, p=0.294 for perimeter and $t_{10}$=1.257, p=0.237 for curvature.

| | Saline | Cocaine-SA |
|---|---|---|
| **Density of synapses** | 2.52<br>synapses / µm$^3$ | No differences (≠) |
| **Proportion of AS:SS** | 96.25:3.75 | No ≠ |
| **Synaptic spatial distribution** | Nearly random | No ≠ |
| **Postsynaptic targets** | The most common synaptic type is axospinous AS (94.21%)<br><br>AS have a preference for dendritic spines (98.34%) and SS for dendritic shafts (82.47%)<br><br>SS are more frequent in spiny shafts than aspiny (94.3 vs 5.7%) | • ≠ for AS:<br>↓ Proportion of AS on spine heads (reduction of 0.7%)<br>↑ Proportion of AS on aspiny shafts (increase of 53%)<br>• No ≠ for SS |
| **Synaptic shape** | The most common synaptic shape is macular both for AS (88.23%) and SS (86.51) | • ≠ for AS:<br>↓ Proportion of macular AS (reduction of 1.8%)<br>↑ Proportion of fragmented AS (increase of 140%)<br>(These ≠ are within axospinous AS but not axodendritic AS)<br>• ≠ for SS:<br>↓ Proportion of macular SS (reduction of 14%)<br>↑ Proportion of horseshoe-shaped SS (increase of 166%)<br>(These ≠ are within axodendritic SS but not axospinous SS) |
| **Synaptic size (SAS area and perimeter)** | AS are smaller than SS (area: 42022 nm$^2$ vs 67972 nm$^2$; perimeter: 996 nm vs 1507 nm)<br><br>Regarding postsynaptic target:<br>Axospinous AS = Axodendritic AS<br>Axospinous SS are smaller than axodendritic SS<br>Synapses on spiny shafts are smaller than on aspiny shafts<br><br>Regarding synaptic shape:<br>Macular AS are smaller than horseshoe-shaped AS (only perimeter), perforated and fragmented synapses<br>Macular SS are smaller than horseshoe-shaped SS | • No ≠ in mean SAS area or perimeter (only ↑ mean SAS perimeter of SS)<br>• ≠ in SAS area or perimeter distribution:<br>For AS:<br>↑ Proportion of larger synapses (All AS, axospinous AS and macular AS)<br>For SS:<br>↑ Proportion of larger synapses (All SS, and tendency for increased proportion of axodendritic SS and perforated SS) |
| **Synaptic curvature (SAS curvature)** | SS are flatter than AS<br><br>No ≠ regarding postsynaptic targets or synaptic shape | No ≠ |
| **Volume fraction of mitochondria** | 7.31% | No ≠ |

**Supplementary table 8. Summary of the main characteristics of normal synaptic organization in the *stratum radiatum* of the rat CA1 and changes after cocaine-SA.** AS: asymmetric synapses; HS: horseshoe-shaped synapses; SA: self-administration; SAS: synaptic apposition surface; SS: symmetric synapses; ≠: differences; ↑: increase; ↓: decrease.

| Group | Spines with one synapse | | Spines with two synapses | | |
|---|---|---|---|---|---|
| | 1 AS | 1 SS | 2 AS | 1 AS + 1 SS | 2 SS |
| Saline | 99.28% (4698) | 0 | 0.11% (5) | 0.61% (29) | 0 |
| Cocaine-SA | 99.37% (4394) | 0 | 0.11% (5) | 0.52% (23) | 0 |

**Supplementary table 9. Single and multiple synapses on dendritic spines in the *stratum radiatum* of the rat CA1 in the saline and cocaine-SA groups.** Data are expressed as percentages and the absolute numbers of synapses studied are given in parentheses. AS: asymmetric synapses; SS: symmetric synapses
